# Identification of Novel Allosteric Sites of SARS-CoV-2 Papain-Like Protease (PLpro) for the Development of COVID-19 Antivirals

**DOI:** 10.1101/2023.05.16.540953

**Authors:** Juliana C. Ferreira, Adrian J. Villanueva, Kenana Al Adem, Samar Fadl, Lara Alzyoud, Mohammad A Ghattas, Wael M. Rabeh

## Abstract

Coronaviruses such as SARS-CoV-2 encode a conserved papain-like protease (PLpro) that is crucial for viral replication and immune evasion, making it a prime target for antiviral drug development. In this study, three surface pockets on SARS-CoV-2 PLpro that may function as sites for allosteric inhibition were computationally identified. To evaluate the effects of these pockets on proteolytic activity, 52 residues were separately mutated to alanine. In Pocket 1, located between the Ubl and thumb domains, the introduction of alanine at T10, D12, T54, Y72, or Y83 reduced PLpro activity to <12% of that of WT. In Pocket 2, situated at the interface of the thumb, fingers, and palm domains, Q237A, S239A, H275A, and S278A inactivated PLpro. Finally, introducing alanine at five residues in Pocket 3, between the fingers and palm domains, inactivated PLpro: S212, Y213, Y251, K254, and Y305. Pocket 1 has a higher druggability score than Pockets 2 and 3. MD simulations showed that interactions within and between domains play critical roles in PLpro activity and thermal stability. The essential residues in Pockets 1 and 2 participate in a combination of intra- and inter-domain interactions. By contrast, the essential residues in Pocket 3 predominantly participate in inter-domain interactions. The most promising targets for therapeutic development are Pockets 1 and 3, which have the highest druggability score and the largest number of essential residues, respectively. Non-competitive inhibitors targeting these pockets may be antiviral agents against COVID-19 and related coronaviruses.

## Introduction

From 2019 to 2022, successive severe acute respiratory syndrome-coronavirus 2 (SARS-CoV-2) variants resulted in global waves of COVID-19 infection ^1^. Although the global incidence rate of COVID-19 has since declined, the race to develop antivirals to combat the impending threat of future viral mutations and antiviral resistance continues. SARS-CoV-2 is a single-stranded RNA virus characterized by crown-like projections on its surface ^2, 3^. The viral genome encodes four structural proteins: spike (S), envelope (E), membrane (M), and nucleocapsid (N) ^4^. Entry of SARS-CoV-2 into the host cell is initiated by binding of the S1 subunit of the S protein to the main host cell receptor, angiotensin-converting enzyme 2 (ACE2) ^5, 6^. The virus then enters the cell via a membrane-fusion step involving the S2 subunit of the S protein ^7^. Upon entry, the host cell machinery translates the viral genomic RNA into two polyproteins, pp1a and pp1ab, which are encoded by the open reading frames (ORFs) ORF1a and ORF1b, respectively. Subsequent cleavage of the polyproteins by two conserved coronaviral proteases produces nonstructural polypeptides (nsps) ^8, 9^. Papain-like protease (PLpro) is required for the proteolytic release of nsps 1–4, whereas 3C-like protease (3CLpro) is required for the proteolytic release of nsps 5–16^10, 11^.

In addition to playing critical roles in viral transcription and replication, PLpro functions in evasion of the host innate immune system. PLpro inhibits the phosphorylation and nuclear translocation of interferon regulatory factor-3 (IRF3), resulting in deactivation of the type I interferon (IFN) response ^12^. Moreover, PLpro is homologous to human deubiquitinases (DUBs) and cleaves ubiquitin and ubiquitin-like modifiers such as interferon-stimulated gene 15 (ISG15). ISGylation and ubiquitination are key elements in the activation of the innate immune response; thus, PLpro inhibits inflammation and antiviral signaling by the immune system and the host’s ability to recognize and degrade invading viral proteins ^12, 13^. Studying the catalytic function and structural stability of PLpro may provide a foundation for developing novel drugs that target viral infection and the immune evasion machinery of multiple coronaviruses, such as SARS-CoV-2, SARS-CoV, and Middle East respiratory syndrome (MERS)-CoV. ^14, 15^

Structural and mutagenesis studies have provided insights into the domains and residues responsible for the catalytic function and stability of PLpro^16–18^. The tertiary structure of the SARS-CoV-2 PLpro monomer adopts a conformation resembling an extended right-handed architecture and comprises two main domains: the ubiquitin-like (Ubl) domain and the catalytic domain ^16–19^. The N-terminal Ubl domain of PLpro consists of residues 1–60, which form a five-stranded β-sheet, an α-helix, and a 3_10_ helix. The catalytic domain comprises residues 61–315, which are divided into the thumb, fingers, and palm domains. The thumb domain (61–176) forms six α-helices and two small β-strands, while the fingers domain (177–238) contains the zinc-binding site and forms a fiveLJstranded βLJsheet and two α-helices. The palm domain (239–315) consists of a sixLJstranded βLJsheet. The catalytic site of PLpro is located between the thumb and palm domains and contains the "catalytic triad" (Cys111, His272, and Asp286), which is essential for the proteolytic mechanism of PLpro ^13, 16, 18^. The Ubl domain binds ubiquitin or ISG15 in an "open hand" structure, with ubiquitin resting on the palm domain. The zinc-binding site, which consists of four conserved cysteine residues (C189, C192, C224, and C226), holds the ubiquitin in place and is essential for the structural integrity and activity of PLpro ^20^. Alanine substitution of these cysteine residues results in complete loss of PLpro function ^16, 21–23^.

Early efforts to develop COVID-19 treatments focused on repurposing approved or investigational drugs against SARS-CoV-2 druggable targets ^24–27^, including PLpro. Many putative candidates against SARS-CoV-2 PLpro identified by *in silico* studies have yet to be biochemically confirmed ^13, 28–31^. Several repurposed protease inhibitors with promising effects on PLpro *in vivo* or *in vitro*, such as simeprevir, tanshinones, famotidine, and ebselen, are either clinically ineffective or still in clinical trials ^32–36^. Another class of inhibitors of interest against PLpro is FDA-approved zinc-ejecting drugs^15, 17, 37, 38^. These drugs target the zinc-finger motif common to metalloenzymes such as PLpro and displace zinc from the zinc-cysteine tetrahedral complex. Although molecular dynamics (MD) simulations of PLpro indicate that the labile zinc-finger domain is highly mobile and thus may be a poor target for small-molecule drugs ^39^, studies have reported that zinc-ejecting drugs such as ebselen and disulfiram reduce the activity of PLpro by disrupting its stability. These findings suggest that zinc-ejecting drugs might be combined with other antiviral drugs to treat COVID-19 symptoms in a so-called "multi-target approach" ^32, 38, 40, 41^.

An alternative to drug repurposing is to identify novel inhibitors of PLpro by computational screening of small-molecule libraries^42–45^. Computational screening typically targets the active site, but this approach is likely to yield competitive inhibitors. Because competitive inhibitors can be outcompeted by peptide substrate, they are considered weaker binders than non-competitive inhibitors, which bind at allosteric sites and are less susceptible to displacement by substrate. The objective of the present study is to characterize surface pockets on PLpro to identify potential allosteric sites for screening small-molecule inhibitors of PLpro.

*In silico* methods were used to identify three novel binding pockets on the surface of SARS-CoV-2 PLpro. Then, site-directed mutagenesis of selected residues in each pocket was performed to determine whether their intra- and inter-domain interactions are important for PLpro activity. Biochemical and thermodynamic analyses of the mutant proteins were conducted, and the experimental data were used to evaluate the druggability of the surface binding pockets. These pockets can be further used in computational screens of small-molecule libraries to identify strong binders that may function as allosteric inhibitors of PLpro and therapeutic agents against COVID-19.

## Results

### Structural assessment of putative binding pockets on the surface of SARS-CoV-2 PLpro

The main objective of this study was to identify potential druggable pockets on the surface of PLpro that may allosterically regulate enzyme activity. Such pockets can be targeted for screening, identifying, and developing new drugs for combating COVID-19. The structure of the PLpro monomer comprises an independent N-terminal ubiquitin-like (Ubl) domain followed by a catalytic region that is composed of an extended right-handed scaffold of three domains: thumb, fingers, and palm (**Figure 1**). The SiteMap module identified four potential druggable pockets on the surface of SARS-CoV-2 PLpro (**Table S1**): the active site and, on the opposite face of the monomer, three potential allosteric binding sites. Pocket 1 is located at the Ubl–thumb interface, Pocket 2 at the thumb–palm–fingers interface, and Pocket 3 at the fingers–palm interface (**Figure 1)**.

**Figure 1.**
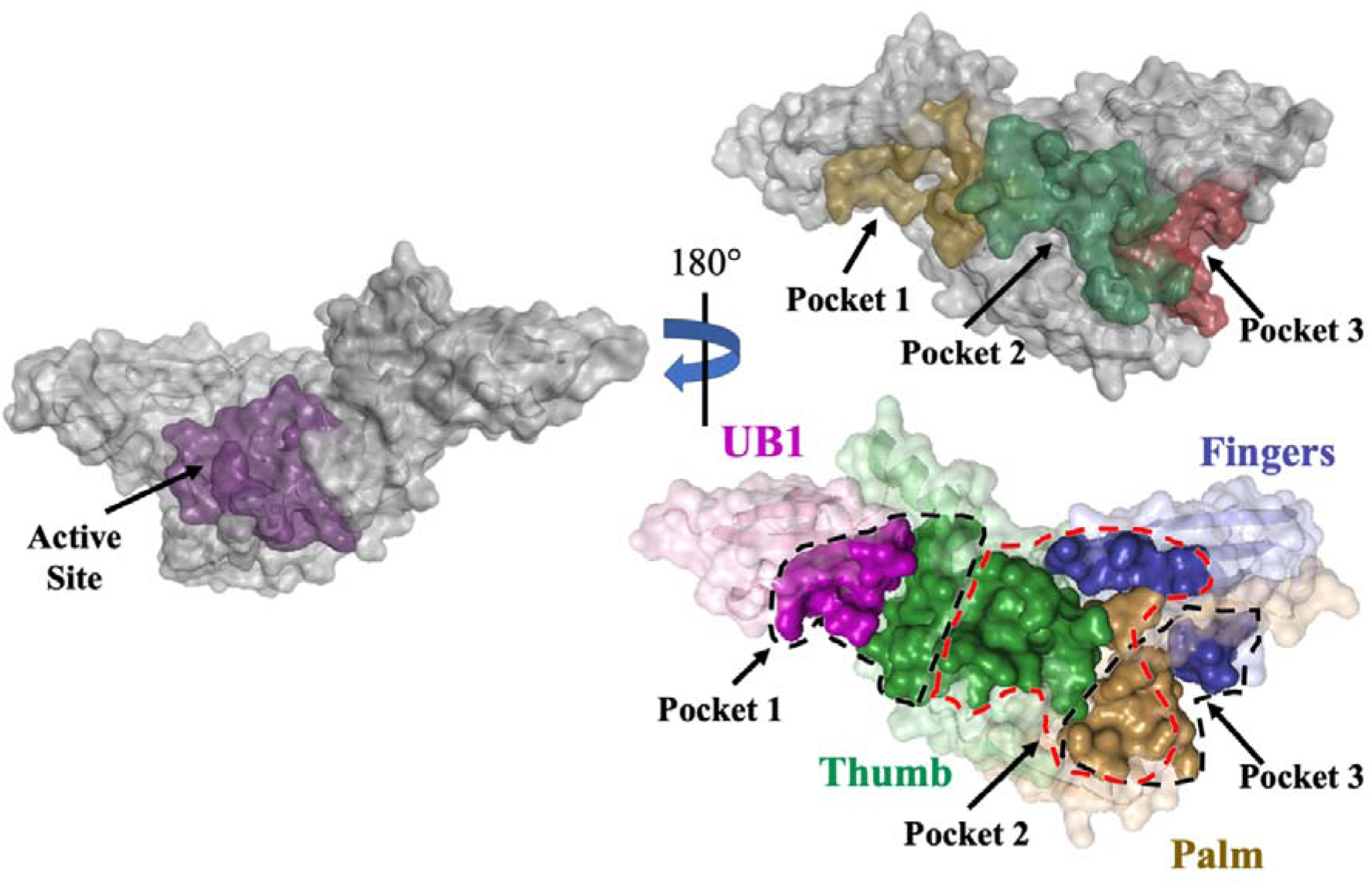
Surface representation of the computationally predicted surface pockets of PLpro. The three pockets are located on the other face of PLpro, opposite the active site. Pocket 1 is a cleft between the thumb and Ubl domains. Pocket 2 is a cavity between the palm, fingers, and thumb domains. Pocket 3 is a cleft between the palm and fingers domains. The PLpro domains are colored as follows: purple, Ubl (residues 1–60); green, thumb (residues 61–176); blue, fingers (residues 177–238); and brown, palm (residues 239–315). The figure was produced using PyMOL Molecular Graphics System version 2.5.5 (Schrödinger LLC).

The active site has the highest druggability score (*D*score), supporting the validity of the computational screen. This high *D*score is the product of a large (*n* = 75 spheres) defined (*e* = 0.68) cavity with high hydrophilicity (*p* = 0.99). The *D*scores of the potential allosteric binding sites illustrate their distinctive physiochemical parameters (**Figure 1** and **Table S1)**. Pocket 1 is large (*n* = 61 spheres) and defined (*e* = 0.67) but has high hydrophilicity (*p* = 1.20), resulting in a *D*score of 0.75. Pockets 2 and 3 are smaller (*n* = 29 and 31 spheres, respectively), less-defined cavities (*e* = 0.58 and 0.62, respectively) with lower hydrophilicity (*p* = 0.89 and 0.99, respectively) than Pocket 1 and have identical *D*scores of 0.57.

The computationally identified pockets encompass a large number of amino acid residues (**Table S2**); thus, site-directed mutagenesis and biochemical analyses were limited to amino acids involved in networks of intra- and inter-domain interactions that may play critical roles in SARS-CoV-2 PLpro activity. To identify potential key residues, the interactions of polar side chains in Pockets 1–3 with other residues were evaluated (**Figures 2–4**). A total of 52 residues were mutated individually to alanine to determine their effects on PLpro activity and thermodynamic stability. The WT and mutant enzymes were expressed in *E. coli* and purified to >90% based on SDS-PAGE analysis. The proteolytic rates of the mutant and WT enzymes were measured at varying enzyme concentrations from 0.5 to 5.0 µM in the presence of a constant peptide substrate concentration of 200 µM. The relative activity of the mutants was calculated by comparing the slope of the plot of proteolytic rate as a function of enzyme concentration with the slope for WT PLpro (**Figures S1–S3**). The alanine mutants were classified into three groups based on their effects on the proteolytic activity of PLpro: active (>60% WT activity), partially active (15%–60% WT activity), and inactive (<15% WT activity). Active mutants were not analyzed further because residues that do not influence activity are not good targets for developing SARS-CoV-2 antivirals.

**Pocket 1.** Of the 18 alanine mutations introduced in Pocket 1, five inactivated and three moderately inactivated PLpro (**Figure 2**). The 18 targeted residues participate in four clusters of interactions. In the first cluster (**Figure 2A**), the side chain hydroxyl of Y71 of the thumb domain forms an intra-domain hydrogen bond with the side chain of D134 at 2.7 Å and an inter-domain hydrogen bond with the backbone of D12 of the Ubl domain at 4.2 Å. In addition, Y72 of the thumb domain forms an inter-domain hydrogen bond with the backbone carbonyl oxygen of V11 of the Ubl domain at 2.6 Å. D134A was fully active, but D12A (8% ± 1% activity) and Y72A were inactive, and Y71A (48% ± 3%) was partially active (**Figure 2E**). In the second cluster (**Figure 2B**), the side chain of D12 of the Ubl domain forms polar intra-domain interactions with the side chains of T10 and N15 at 3.4 Å and 2.8 Å, respectively, and the backbone of N15 at 2.7 Å. Additionally, E67 of the thumb domain forms two polar interactions with the Ubl domain: the imidazole moiety of H17 at 3.4 Å and the carboxamide moiety of N15 at 4.4 Å. Like D12A, T10A was inactive; E67A (18% ± 1%) was partially active, and N15A and H17A (77% ± 1%) were active (**Figure 2E**). In the third cluster (**Figure 2C**), K91 of the thumb domain forms two polar interactions: an intra-domain interaction with N88 at 2.9 Å and an inter-domain interaction with D37 of the Ubl domain at 4.6 Å. However, K91A, N88A, and D37A were all active (**Figure 2E**). Finally, in the fourth cluster (**Figure 2D**), the side chain of N13 of the Ubl domain forms two intra-domain interactions with the hydroxyl groups of T54 and Y56 at 4.7 Å and 4.2 Å, respectively. In addition, the side chain of Y56 of the Ubl domain may form a second hydrogen bond with the side chain of Y83 of the thumb domain at 4.5 Å. The guanidine group of R138 of the thumb domain forms two intra-domain bonding interactions: a salt bridge with E143 at 3.1 Å and a hydrogen bond with N146 at 3.4 Å, and N146 forms a tight hydrogen bond with Y83 at 2.6 Å. Interestingly, N13A was active, whereas T54A was inactive and Y56A was partially active (44% ± 1%) (**Figure 2E**). Y83A was inactive (12% ± 1%), but R138A, E143A, and N146A were all fully active (**Figure 2E**).

**Figure 2.**
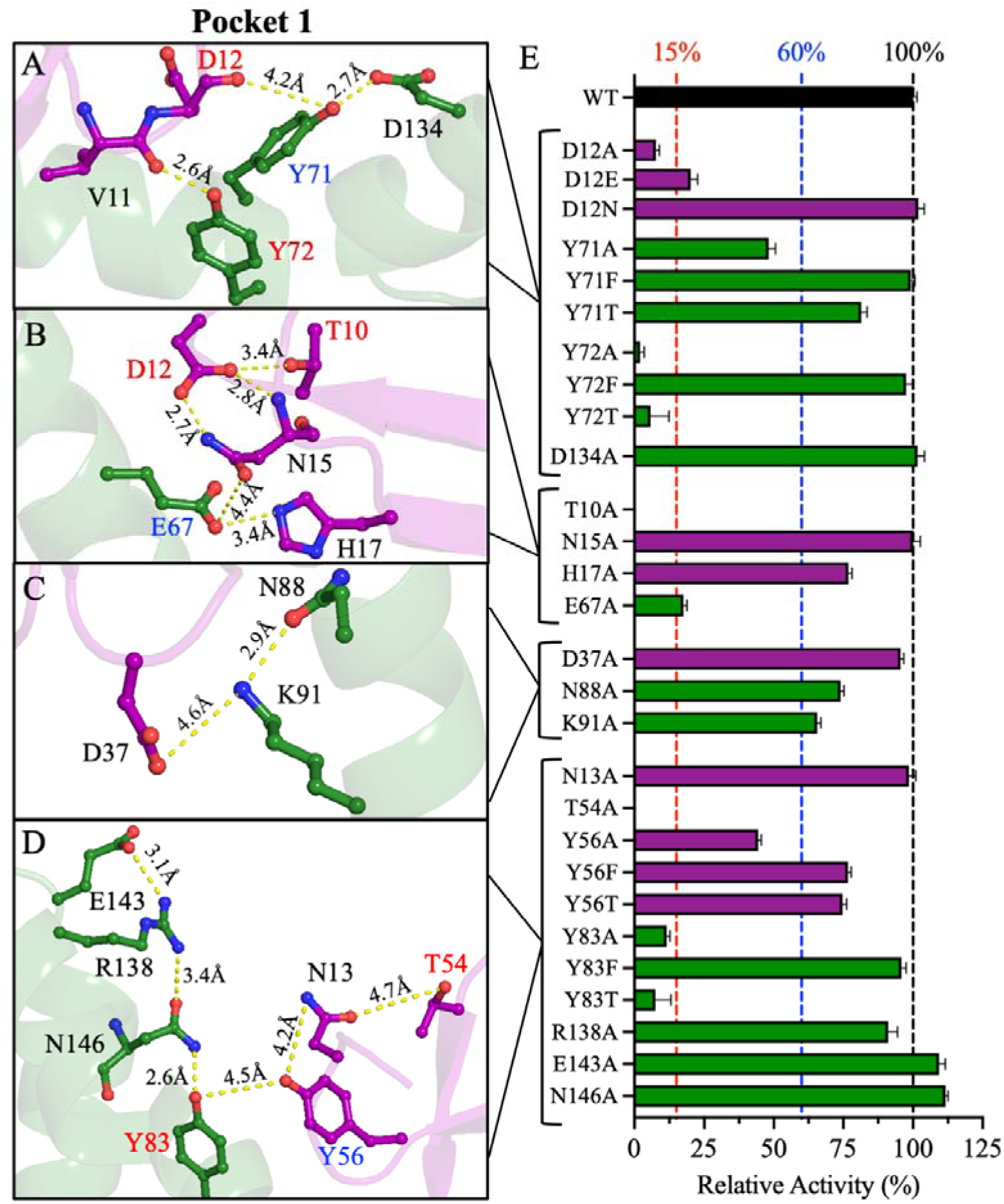
Pocket 1 of PLpro: structural analyses and relative activities. (**A**–**D**) Structural representations of key residues in Pocket 1 at the interface of the Ubl and thumb domains. Only residues participating in side chain–side chain or side chain–backbone interactions are considered. The labels of the residues are color-coded based on their effects on the relative rate of PLpro upon site-directed mutagenesis: red for mutants with <15% relative activity, blue for mutants with relative activity of 15%–60%, and black for mutants with >60% relative activity. The figure was produced using PyMOL Molecular Graphics System version 2.5.5 (Schrödinger LLC). (**E**) Bar plot of the relative activities of Pocket 1 mutants compared to WT. The relative activities were calculated from the enzyme titration data in Figure S1. The colors of the residues in A–D and the bars in E correspond to the domain colors established in Figure 1 (purple = Ubl and green = thumb). The data are presented as the mean ± SD, n=3.

To further investigate the roles of the residues whose mutation to alanine significantly reduced PLpro activity, additional mutants were constructed in which the original amino acid was replaced with an amino acid with similar physicochemical properties. In the first cluster of Pocket 1, the D12E mutation partially recovered activity compared to the D12A mutation, whereas D12N had full WT activity (**Figure 2E**). Similarly, Y71T and Y71F partially and completely recovered WT activity, respectively. Interestingly, while Y72F had full WT activity, Y72T was inactive, despite potentially maintaining polar side chain interactions with the carbonyl oxygen of the backbone of V11. In the fourth cluster, Y56 of the Ubl domain and Y83 of the thumb domain facilitate inter-domain interactions. Y56T and Y56F partially recovered PLpro activity. Surprisingly, Y83T failed to restore proteolytic activity despite retaining a polar hydroxyl side chain, whereas Y83F fully restored PLpro activity. These data indicate that the hydrophobic interactions initiated by some residues of Pocket 1 are more crucial for activity than their polar interactions are (**Figure 2E**).

**Pocket 2.** Pocket 2 is unique among the potential allosteric sites because it lies at the interface of the thumb, fingers, and palm domains of the catalytic region (**Figure 1**). Site-directed mutations in this pocket targeted four clusters of interactions. In the first cluster, which lies at the palm– thumb interface (**Figure 3A**), three inter-domain hydrogen bonds are possible between the side chain of H275 of the palm domain and the side chains of T115, T119, and Q122 of the thumb domain at 3.0 Å, 3.6 Å, and 2.8 Å, respectively. Of the four alanine mutants, only H275A was inactive (**Figure 3E**). Q122A was partially active (35% ± 1%), and Q122E partially restored activity (52% ± 3%). T115A was active (79% ± 1%), but surprisingly, T119A had higher relative activity than WT (141% ± 1%). Q122 of the thumb domain also forms an inter-domain hydrogen bond with the side chain hydroxyl of T277 of the palm domain at 2.7 Å (**Figure 3A**). In addition, T277 may form a long-range intra-domain hydrogen bond with the side chain of K279 at 5.1 Å. T277A was partially active (32% ± 3%), whereas K279A was fully active (**Figure 3E**). Introducing T277S rescued proteolytic activity (81% ± 1%). K279 also participates in a set of intra-domain interactions involving H255, G256, and S278 of the palm domain (**Figure 3A**). The side chain of S278 may form three possible hydrogen bonds with the backbone amide groups of G256 and K279 at 2.9 Å and 3.7 Å, respectively, and with the side-chain imidazole group of H255 at 4.2 Å. H255A and S278A were partially active (43% ± 2% and 16% ± 1%, respectively), whereas K279A was fully active (**Figure 3E**). Despite the structural similarity between serine and threonine, introducing S278T did not restore activity (7% ± 4%).

**Figure 3.**
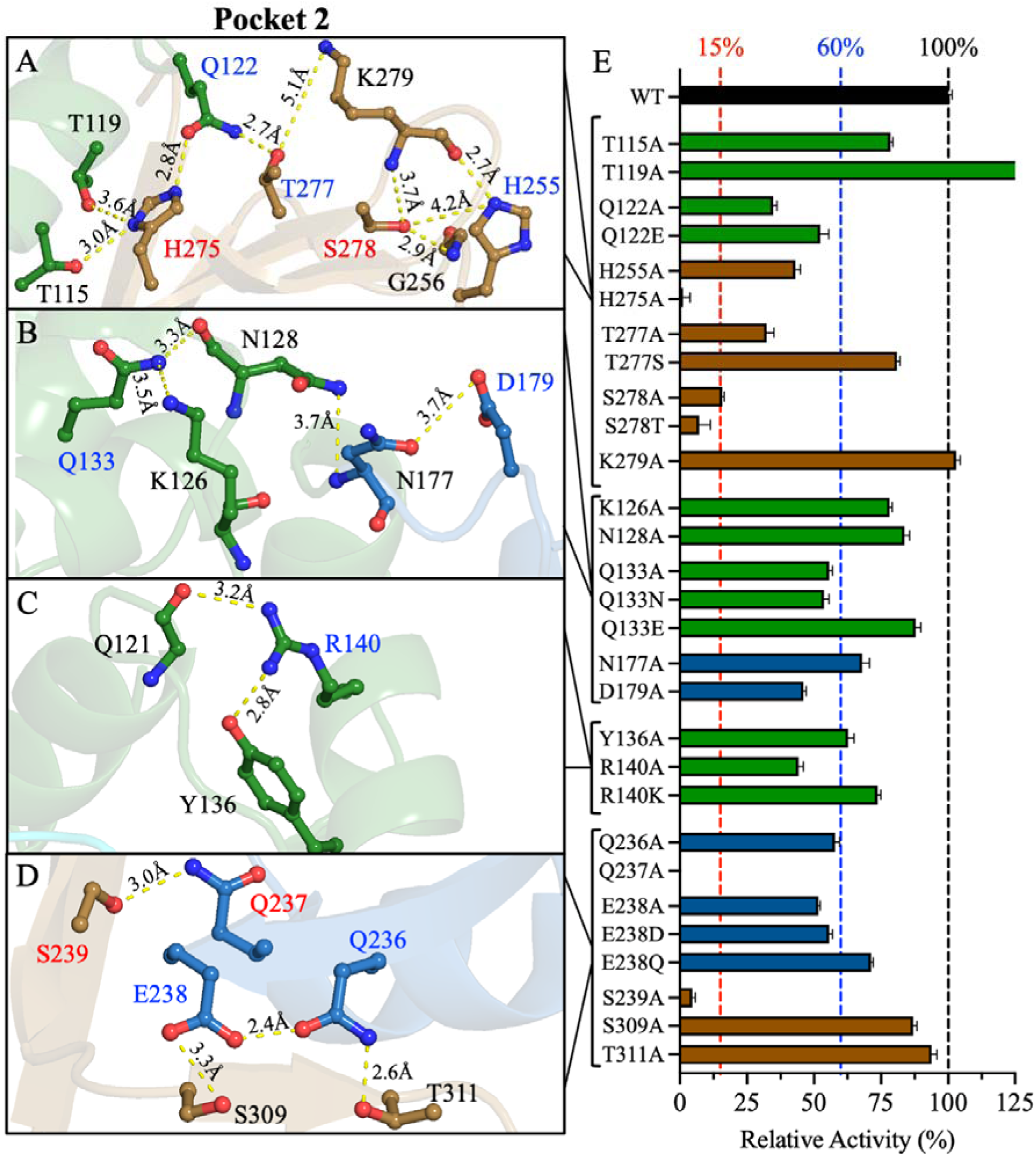
Pocket 2 of PLpro: structural analyses and relative activities. (**A**–**D**) Structural representations of key residues in Pocket 2 at the interface between the thumb, fingers, and palm domains. The residue labels are color-coded as described in Figure 2. The figure was produced using PyMOL Molecular Graphics System version 2.5.5 (Schrödinger LLC). (**E**) Bar plot of the relative activities of Pocket 2 mutants compared to WT. The relative activities were calculated from the enzyme titration data in Figure S1. The colors of the residues in A– D and the bars in E correspond to the domain colors established in Figure 1 (green = thumb, blue = fingers, and brown = palm). The data are presented as the mean ± SD, n=3.

In the second cluster, polar interactions were observed at the thumb–fingers interface (**Figure 3B**). The side chain of Q133 of the thumb domain initiates multiple intra-domain interactions, including two possible hydrogen bonds with the side chain of K126 at 3.5 Å and the backbone of N128 at 3.3 Å. Q133A was partially active (56% ± 1%), whereas K126A and N128A were active (78% ± 1% and 84% ± 2%, respectively) (**Figure 3E**). Introducing Q133N did not rescue the activity of PLpro, but Q133E restored activity (88% ± 2%). In the fingers domain, the side chains of N177 and D179 form an intra-domain hydrogen bond at 3.7 Å (**Figure 3B**). The only inter-domain interaction is between the side chain of N128 of the thumb domain and the side chain of N177 of the fingers domain at 3.7 Å. N177A (68% ± 3%) and D179A (46% ± 1%) were partially active (**Figure 3E**).

The third cluster lies within the thumb domain and features short interactions initiated by the side chain of R140, which forms dual hydrogen bonds with the side chain hydroxyl of Y136 at 2.8 Å and the backbone carbonyl oxygen of Q121 at 3.2 Å (**Figure 3C**). R140A was partially active (44% ± 2%), whereas Y136A remained active (63% ± 2%) (**Figure 3E**). Introducing R140K recovered activity to 74% ± 1%.

The final set of interactions in Pocket 2 lies at the fingers–palm interface (**Figure 3D**). The side chain of Q237 forms an inter-domain interaction with the side chain hydroxyl of S239 at 3.0 Å, and both Q237A and S239A were inactive (**Figure 3E**). The side chains of Q236 and E238 of the fingers domain form inter-domain interactions with the side chains of T311 and S309 of the palm domain at 2.6 Å and 3.3 Å, respectively (**Figure 3D**). The only intra-domain interaction in this cluster is a hydrogen bond between the side chains of Q236 and E238 at 2.4 Å. S309A and T311A remained active (**Figure 3E**), whereas Q236A and E238A were partially active (58% ± 2% and 52% ± 1%, respectively). E238D was also partially active (56% ± 1%), but introducing E238Q restored activity (71% ± 1%).

**Pocket 3.** Pocket 3 is located at the fingers–palm interface and features multiple inter-domain interactions between S212 and Y213 of the fingers domain and Y251, Y305, and E307 of the palm domain (**Figure 4B**). Hydrogen bonds are formed between the side chains of S212 and Y251 at 4.4 Å, the side chains of Y213 and E307 at 2.5 Å, and the side chain hydroxyl group of Y305 and the backbone amide group of Y213 at 3.5 Å. The only intra-domain interactions in this group of residues are between the side chains of K217 and E214 of the fingers domain at 3.2 Å and between the side chains of Y251 and Y305 of the palm domain at 4.1 Å. All alanine mutations of tyrosine residues (Y213A, Y251A, and Y305A) completely inactivated PLpro, and also S212A was classified as inactive (9% ± 6%) (**Figure 4A**). E214A, K217A, and E307A were partially active (49% ± 2%, 21% ± 3%, and 29% ± 4%, respectively) (**Figure 4A**). These data highlight the importance of the Pocket 3 network of interactions for PLpro activity. Introducing the alternative amino acid substitution Y213T failed to rescue PLpro activity, whereas Y213F partially rescued activity (28% ± 2%). On the other hand, introducing Y305F or Y305T did not recover the PLpro activity. By sharp contrast, both Y251F and Y251T were active (91% ± 1% and 90% ± 1%, respectively). In addition, S212T was also inactive as was the case with S212A.

**Figure 4.**
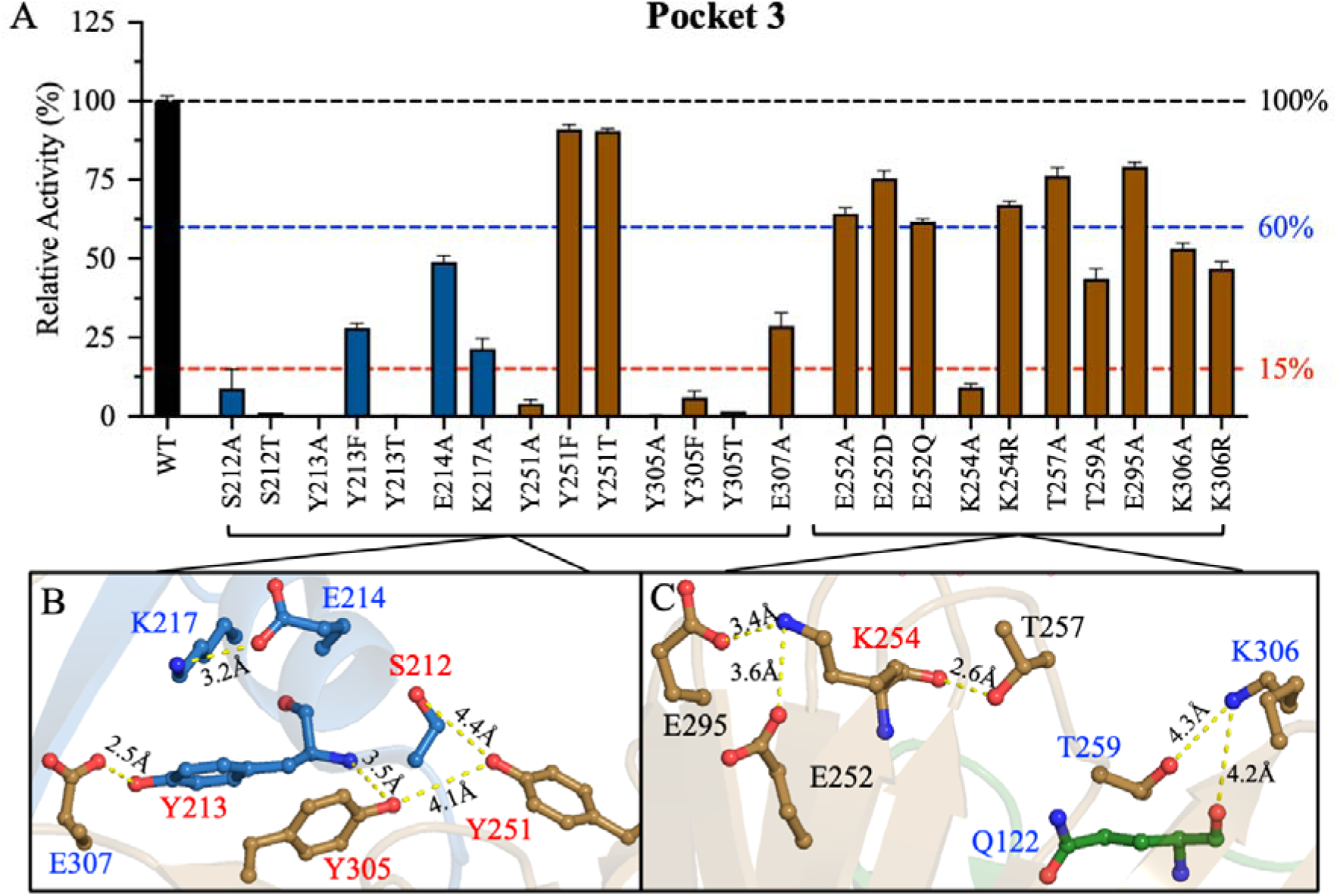
Pocket 3 of PLpro: structural analyses and relative activities. (**A**) Bar plot of the relative activities of Pocket 3 mutants compared to WT. The relative activities were calculated from the enzyme titration data in Figure S1. (**B**–**C**) Structural representations of key residues in Pocket 3, a cavity formed between the fingers and palm domains. The residue labels are color-coded as described in Figure 2. The figure was produced using PyMOL Molecular Graphics System version 2.5.5 (Schrödinger LLC). The colors of the bars in A and the residues in B–C correspond to the domain colors established in Figure 1 (blue = fingers and brown = palm). The data are presented as the mean ± SD, n=3.

Another cluster of interactions in Pocket 3 is located in the palm domain, apart from a single inter-domain interaction with Q122 of the thumb domain (**Figure 4C**). The side chain of K254 may form a salt bridge with the side chain of E252 or E295 at 3.6 Å or 3.4 Å, respectively. Moreover, the backbone of K254 forms an intra-domain hydrogen bond with the side chain hydroxyl of T257 at 2.6 Å. Among the alanine mutations of these residues, only K254A inactivated PLpro (9% ± 1%), and K254R rescued proteolytic activity (67% ± 1%) (**Figure 4A**). E252A, T257A, and E295A remained active (64% ± 2%, 76% ± 3%, and 79% ± 1%, respectively). Compared with E252A, E252Q did not recover activity (62% ± 1%), but E252D restored it slightly (75% ± 3%). Finally, the side chain of K306 forms an intra-domain interaction with the side chain of T259 at 4.3 Å and an inter-domain interaction with the backbone carbonyl oxygen of Q122 at 4.2 Å (**Figure 4C**). Q122 is also involved in interactions in Pocket 2 with T277 and H275 of the palm domain. Q122A, T259A, and K306A were all partially active (35% ± 1%, 44% ± 3%, and 53% ± 2%, respectively) (**Figure 3E and 4E**). Compared with K306A, introducing K306R did not improve PLpro activity (47% ± 2%).

### Initial velocity studies of PLpro mutants

Initial velocity studies were performed to determine the turnover number (*k_cat_*), Michaelis constant (*K*_m_), and catalytic efficiency (*k_cat_*/*K*_m_) of WT PLpro and selected mutants (**Figure 5, Figures S4-S5, and Tables S3-S5**). All partially active PLpro mutants, several of the recovery mutants, and T119A, the only mutant with enhanced activity, were characterized. Some PLpro mutants with very low relative activity were characterized by using high enzyme concentrations in the assays for those mutants. However, mutants classified as inactive in the enzyme titration experiments, such as T10A and Y72A, were not investigated because their activity was below the experimental detection limits.

**Figure 5:**
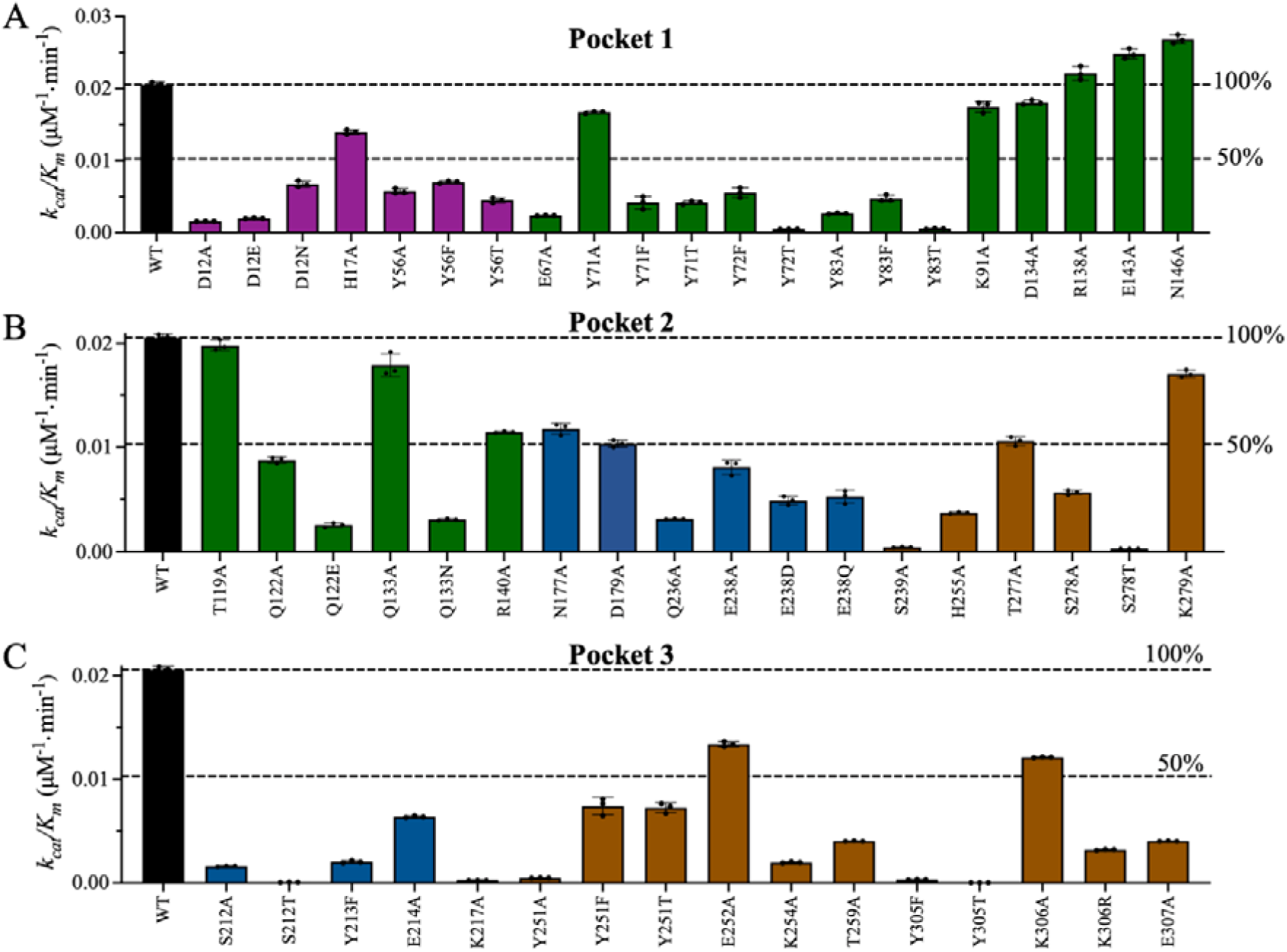
Kinetic parameters of the enzymatically active PLpro mutants. Bar plots of the catalytic efficiency (*k_cat_*/K_m_) of WT PLpro (black bars) **(A)** Pocket 1 mutants, **(B)** Pocket 2 mutants, and **(C)** Pocket 3 mutants. Initial velocity studies were conducted for all mutants with experimentally measurable enzymatic activity. Some mutants for which data are shown had relative activities of less than 8%. The bar plot colors correspond to the domain colors established in Figure 1 (purple = Ubl, green = thumb, blue = fingers, and brown = palm). The data points represent the mean ± SD, n=3.

**Pocket 1.** Y71 and Y72 of the thumb domain interact with the backbones of V11 and D12 of the Ubl domain. Y71A and its rescue mutations Y71F and Y71T decreased *k_cat_* by up to 2.5-fold compared to WT (**Figure S4A and Table S3**). Compared with WT, Y71A decreased *K*_m_ by 1.7-fold, whereas Y71F and Y71T increased *K*_m_ by 3- and 2-fold, respectively (**Figure S5A**). Because the *K*_m_ and *k_cat_* of Y71A were both reduced compared to those of WT, the catalytic efficiency, *k_cat_*/*K*_m_, of Y71A was similar to that of WT. By contrast, Y71F and Y71T both reduced catalytic efficiency by 4.9-fold compared to WT (**Figure 5A and Table S3**). Y71 also interacts with D134; however, the kinetic parameters of D134A were similar to those of WT. Because Y72A had low relative activity (2% of WT), initial velocity studies of this mutant could not be performed, and the rescue mutants, Y72T and Y72F, were evaluated instead. Y72T and Y72F reduced *k_cat_*by 22.0- and 2.1-fold, respectively, compared to WT (**Figure S4A and Table S3**), and increased K_m_ by 1.8-fold (**Figure S5A**). Overall, Y72T and Y72F reduced catalytic efficiency by 40.2- and 3.7-fold, respectively, compared to WT (**Figure 5A and Table S3**). These data indicate that replacing Y72 with phenylalanine (but not threonine) preserved the interaction with the backbone of V11 and PLpro activity.

The initial velocity studies also provided insights into the interactions of D12 and H17 of the Ubl domain with T10, N15, and E67 of the thumb domain. D12A and its rescue mutants D12E and D12N decreased *k_cat_* by 10.8-, 5.9-, and 1.7-fold, respectively, compared to WT (**Figure S4A** and **Table S3**). The *K*_m_ values of the D12 mutants were approximately 1.8-fold higher than that of WT (**Figure S5A** and **Table S3**). Overall, D12A and D12E reduced *k_cat_*/*K*_m_ by ∼10-fold compared to WT, whereas D12N decreased *k_cat_*/*K*_m_ by only 3.1-fold, indicating that the introduction of asparagine preserved interactions with Y71 (**Figure 5A** and **Table S3**). H17A and E67A reduced *k*_cat_ by 1.7- and 6.8-fold, respectively, compared to WT, but altered *K*_m_ only slightly (**Figures S4A** and **S5A**). Thus, H17A and E67A reduced *k_cat_*/K_m_ by 1.5- and 8.6-fold, respectively, compared to WT (**Figure 5A**).

Finally, in Pocket 1, Y56 of the Ubl domain interacts with Y83 of the thumb domain. Compared to WT, Y56A and its rescue mutations Y56F and Y56T reduced *k_cat_* by 1.7- to 2.8-fold while increasing *K*_m_ by up to 2-fold (**Figure S4A**, **S5A,** and **Table S3**). As a result, all Y56 mutations reduced *k_cat_*/K_m_ by 2.9- to 4.6-fold compared to WT (**Figure 5A**). Y83A and Y83T reduced *k_cat_* by 8.6- and 19-fold, respectively, and altered *K*_m_ slightly, resulting in reductions of *k_cat_*/K_m_ of 7.7- and 37.3-fold, respectively, compared to WT (**Figure 5A, S4A, and S5A**). Compared with Y83A, the rescue mutant Y83F exhibited improved kinetic parameters. Y83F reduced *k*_cat_ by 2.2- fold, increased *K*_m_ by 2-fold, and decreased *k_cat_*/K_m_ by 4.3-fold compared to WT. The kinetic parameters of K91A, R138A, E143A, and N146A were not greatly different from those of WT.

**Pocket 2.** Pocket 2 includes multiple residues at the interface of the thumb, fingers, and palm domains. Q122 of the thumb domain interacts with H275 and T277 of the palm domain (**Figure 3A**). Kinetic parameters were not obtained for H275A due to its low relative activity. Q122A Q122E, and T277A decreased *k_cat_* by 2.9- to 3.9-fold compared to WT. Q122A and T277A had minor effects on *K*_m_, whereas Q122E increased *K*_m_ by 2-fold. Overall, Q122A, Q122E, and T277A reduced *k_cat_*/*K*_m_ by 2.4-, 8-, and 1.9-fold compared to WT, respectively (**Figure 5B, Figures S4B and S5B, and Table S4**). These results indicate that the interaction between Q122 and T277 is important for maintaining active PLpro and that glutamate substitution at Q122 is not sufficient to preserve PLpro activity.

Intra-domain interactions of the palm domain in Pocket 2 include interactions of H255 and S278 with the backbone of K279 (**Figure 3A**). Compared to WT, H255A, S278A, and S278T decreased *k_cat_* by 4.3-, 5.3-, and 28.7-fold and *k_cat_*/K_m_ by 5.5-, 3.6-, and 62.7-fold, respectively. The decreases in *k_cat_*/K_m_ reflect the fact that only S278T affected *K*_m_ (2.2-fold increase) (**Figure 5B, S4B, S5B, and Table S4**). The lower catalytic efficiency of S278T compared to S278A suggests that the bulkier size of threonine reduced the rate of catalysis.

Q133, R140, and N177 in Pocket 2 facilitate intra-domain interactions in the thumb domain (**Figure 3B** and **3C**). Alanine substitutions at these residues did not greatly alter the kinetic parameters of PLpro (**Figure 5B, S4B, S5B, and Table S4**). However, Q133N reduced *k*_cat_ by 3.3-fold, increased *K*_m_ by 2-fold, and reduced *k_cat_*/*K*_m_ by 6.6-fold compared to WT. Thus, the catalytic efficiency of Q133N was lower than that of Q133A. D179 also forms intra-domain interactions in the thumb domain, and D179A increased *K*_m_ by 2-fold compared to WT but did not alter *k*_cat_.

Inter-domain interactions in Pocket 2 between Q236, Q237, and E238 of the fingers domain and S239, S309, and T311 of the palm domain are essential for PLpro activity (**Figure 3D**). Q236A, E238A, and S239A reduced *k_cat_*by 4.6-, 1.6-, and 32.6-fold, respectively (**Figures 5B, S4B and S5B**), increased *K*_m_ slightly, and reduced *k_cat_*/K_m_ by 6.5-, 2.5-, and 46.9-fold compared to WT. The rescue mutations E238D and E238Q had similar effects on the kinetic parameters of PLpro, including a small decrease in *k_cat_*, a slight increase in *K*_m_, and ∼4-fold reduction in *k_cat_*/*K*_m_. The kinetic parameters of Q237A were not acquired due to its low relative activity.

**Pocket 3.** Nearly all mutations in Pocket 3 reduced catalytic efficiency by <50% relative to WT. S212, Y213, E214, and K217 of the fingers domain participate in inter-domain interactions with Y251, Y305, and E307 of the palm domain (**Figure 4B**). Initial velocity studies were conducted with S212A, E214A, K217A, Y251A, and E307, which had sufficient catalytic rates for measurement experimentally. The phenylalanine and threonine mutants of Y213 and Y305 were also characterized. S212A, E214A, K217A, Y251A, and E307A reduced *k_cat_* by 9.6-, 1.7-, 36.5-, 31.9-, and 4.1-fold, respectively, increased *K_m_* by 1.3- to 1.9-fold, and decreased *k_cat_*/*K_m_* by 12.7-, 3.2-, 69.7-, 2.8-, and 5.1-fold compared to WT, respectively (**Figures 5C, S4C, and S5C and Table S5**). The alternative substitutions Y213F, Y305F, and Y305T decreased *k_cat_* by 5.5-, 19.6-, and 297.1-fold, increased *K_m_*by 1.9-, 3.3-, and 2.6-fold, and reduced *k_cat_*/*K_m_*by 10.2-, 64.3-, and 755.6-fold, respectively, compared to WT. Compared with WT, S212T reduced *k_cat_* by 126.3- fold and thus did not rescue S212A. By contrast, Y251F and Y251T reduced *k_cat_*/*K_m_*by only 2.8- fold compared to WT, whereas Y251A reduced *k_cat_*/*K_m_*by 42.4-fold (**Figure 5C** and **Table S5**).

Intra-domain interactions occur between E252, K254, T259, and K306 of the palm domain (**Figure 4C**). The kinetic parameters of E252A were similar to those of WT. By contrast, K254A, T259A, and K306A reduced *k_cat_* by 10.8-, 4.0- and 3.3-fold and *k_cat_*/*K_m_* by 10.5-, 5.0- and 6.4-fold compared to WT (**Figure 5C, S4C, S5C, and Table S5**). The rescue mutant K306R had lower catalytic efficiency than K306A, which may indicate that the size of the side chain at this position is important for PLpro activity (**Figure 5C**).

### Thermodynamic and kinetic stabilities of the PLpro mutants

To investigate the correlation between the activity and thermodynamic stability of PLpro, differential scanning calorimetry (DSC) of WT and mutant PLpro was performed. The PLpro thermograms displayed two-state transitions. The melting point (*T*_m_) was determined from the apex of the peaks, and the calorimetric enthalpy (Δ*H*_cal_) was calculated from the area under the thermographic peaks (**Figure S6**). WT PLpro displayed two melting points, T_m1_ and T_m2_, at 44 °C and 52 °C, respectively, and Δ*H*_cal_ of 1094 kJ/mol. Similar to WT PLpro, all inactive mutants displayed two distinct melting points. Compared with WT, *T*_m1_ was higher for all inactive mutants except S278A (**Figure S6D, S6E** and **Table S6**). Similarly, all inactive mutants except K254A exhibited an increase in *T*_m2_ of up to 6 °C compared with WT (**Figure S6E**). K254A in Pocket 3 decreased *T*_m2_ by 1.5 °C compared to WT. Moreover, most of the inactive mutants exhibited considerable reductions in Δ*H*_cal_ compared to WT (**Figure S6F** and **Table S6**), implying that these mutations altered the structural fold of PLpro. The exceptions were Y213A in Pocket 3, which increased Δ*H*_cal_ compared to WT (1.7-fold), and S239A, which resulted in Δ*H*_cal_ similar to that of WT.

Thermal inactivation kinetics studies of WT PLpro and the partially active mutants were performed to elucidate the relationship between structural stability and activity (**Figure 6**). PLpro was incubated at 37 °C, and the residual enzymatic activity was determined at different time points for up to two hours. The kinetic half-life (*t*_1/2_) is the time at which 50% relative activity remains and was used as a quantitative measure of the thermal stability of PLpro variants. Inactive mutants were not included in these experiments because of the difficulty of reliably calculating *t*_1/2_. The *t*_1/2_ of WT PLpro was 91 min (**Figure 6A and 6G**). In Pocket 1, the partially active mutants—Y56A, E67A, and Y71A—had longer *t*_1/2_ values of 133 min, 125 min, and 185 min, respectively (**Figure 6A** and **6G**), indicating greater thermal stability. Most notably, the *t*_1/2_ of Y71A was ∼2-fold higher than that of WT. By contrast, some mutations in Pocket 2 reduced thermal stability compared to WT (**Figure 6B–D** and **6G**). Q122A, E238A, T277A, and S278A had reduced *t*_1/2_ values of 12 min, 77 min, 56 min, and 11 min, respectively.

**Figure 6.**
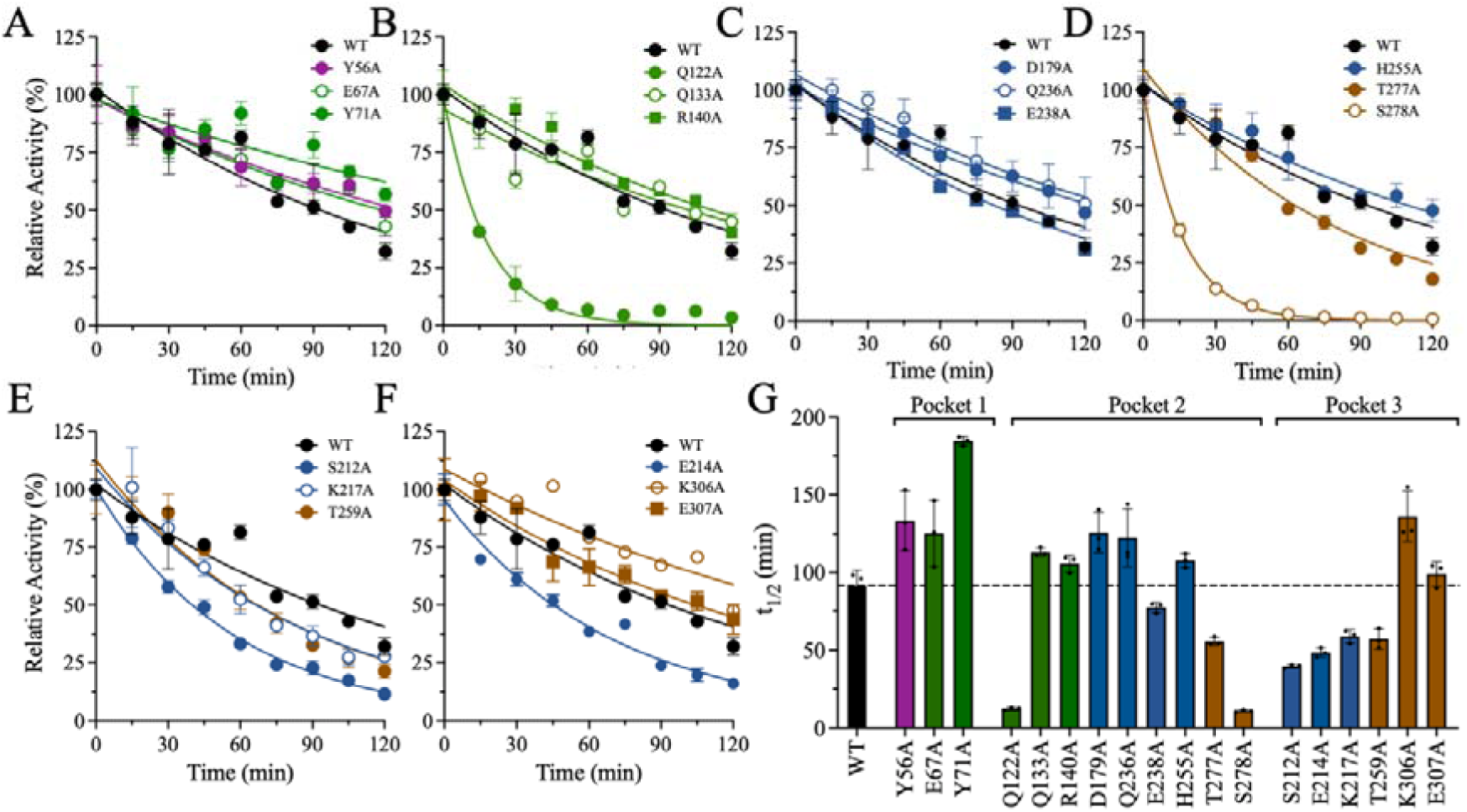
Thermal inactivation kinetic measurements and half-life (t_1/2_) calculation. The residual enzymatic activity of each PLpro variant was measured at 15-minute intervals while incubating the enzyme at 37 °C for up to two hours: (**A**) Pocket 1 mutants, (**B**–**D**) Pocket 2 mutants, and (**E**–**F**) Pocket 3 mutants. **(G)** Bar plot of the half-life values of WT and mutant PLpro. The bar plot colors correspond to the domain colors established in Figure 1 (purple = Ubl, green = thumb, blue = fingers, and brown = palm). The data points represent the mean ± SD, n=3.

Among all mutations in all pockets, Q122A and S278A resulted in the largest decreases in t_1/2_, with reductions of 7- to 8-fold compared to WT. The other mutations in Pocket 2—Q133A, R140A, D179A, Q236A, and H255A—increased *t*_1/2_ slightly compared to WT. Finally, the Pocket 3 mutations S212A, E214A, K217A, and T259A reduced *t*_1/2_ to 40–60 min. The other mutations in Pocket 3 either did not affect *t*_1/2_, *i.e.,* E307A, or increased *t*_1/2_ to 136 min, *i.e.,* K306A (**Figure 6E–G**).

DSC was performed to further evaluate the thermodynamic stability of the mutants with lower *t*_1/2_ values than WT: Q122A, E214A, K217A, E238A, T259A, T277A, and S278A (**Figure S7** and **Table S6**). Similar to the DSC thermograms of the inactive mutants, the thermograms of the partially active mutants included a two-state transition with two melting points (*T*_m1_ and *T*_m2_). Only S278A decreased *T*_m1_ compared to WT, from 44.4 ± 0.4 °C to 43.7 ± 0.2°C. The other mutations increased *T*_m1_ to values between 45.6 ± 1.0 °C and 50.2 ± 0.4 °C; T259A had the highest *T*_m1_ (**Figure S7C**). Most of the mutations also increased *T*_m2_ from 51.9 ± 0.1 °C for WT to values between 53.2 ± 0.1 °C and 56.7 ± 0.5 °C. Again, T259A had the highest *T*_m2_ (**Figure S7D**). Only two mutants, E238A and T277A, had *T*_m2_ values similar to that of WT. Overall, T259A exhibited the largest increases in T_m1_ and T_m2_ (6 °C and 5 °C, respectively). Finally, nearly all mutations decreased Δ*H*_cal_ compared to WT. Δ*H*_cal_ was 1094 ± 73 kJ/mol for WT but ranged between 618 ± 49 kJ/mol and 921 ± 59 kJ/mol for the mutants (**Figure S7E**). The exception was T277A, which increased Δ*H*_cal_ by 3-fold to 2749 ± 146 kJ/mol (**Table S6**). Collectively, the thermal analyses presented here confirm that the mutant enzymes are thermodynamically stable; thus, the loss of activity is not a result of structural instability. The mutations may have altered the overall structural fold such that the enzyme is less active than WT PLpro but has similar thermodynamic stability.

### Assessment of protein stability by molecular dynamics (MD) simulations

MD simulations were performed to assess the effects of mutations of residues in the three pockets predicted to be essential for the catalytic function of PLpro. Among the mutations in Pockets 1 and 2, D12A, Y72A, T10A, T54A, Y83A, H275A, and S239A reduced the catalytic activity of PLpro but did not alter protein dynamics compared to WT PLpro (data not shown). Pocket 3 contains the most residues that contribute to the catalytic activity of PLpro. MD simulations were conducted on S212A, Y213A, Y251A, K254A, and Y305A, as all of these mutations in Pocket 3 reduced or completely eliminated the catalytic activity of PLpro. Y251A and Y305A in the palm domain exhibited the largest changes in dynamics. The root mean square deviation (RMSD) of the palm domain was examined to elucidate the impact of the Y251A and Y305A mutations on structural and conformational dynamics. Both Y251A and Y305A significantly disturbed the stability of PLpro throughout the 200-ns MD simulation time and increased RMSD compared to WT (**Figure 7A**). The RMSD values of WT fluctuated between 1.5 Å and 2.5 Å throughout the simulations. For Y251A, the RMSD value gradually increased from 1.5 Å and oscillated between 2.5 Å and 3.5 Å, with peaks reaching almost 4 Å at 55 ns and 95 ns. The RMSD of Y305A was also 1.5 Å at the beginning of the simulation and steadily increased until reaching ∼4.5 Å at ∼115 ns; it then briefly dropped to oscillate around ∼2.5 Å before increasing again to fluctuate at ∼3.5 Å. These deviations reflect sustained conformational changes in Y251A and Y305A at various stages of the MD simulation, indicating that these mutations reduce the activity of PLpro by increasing the structural dynamics of the protease.

**Figure 7.**
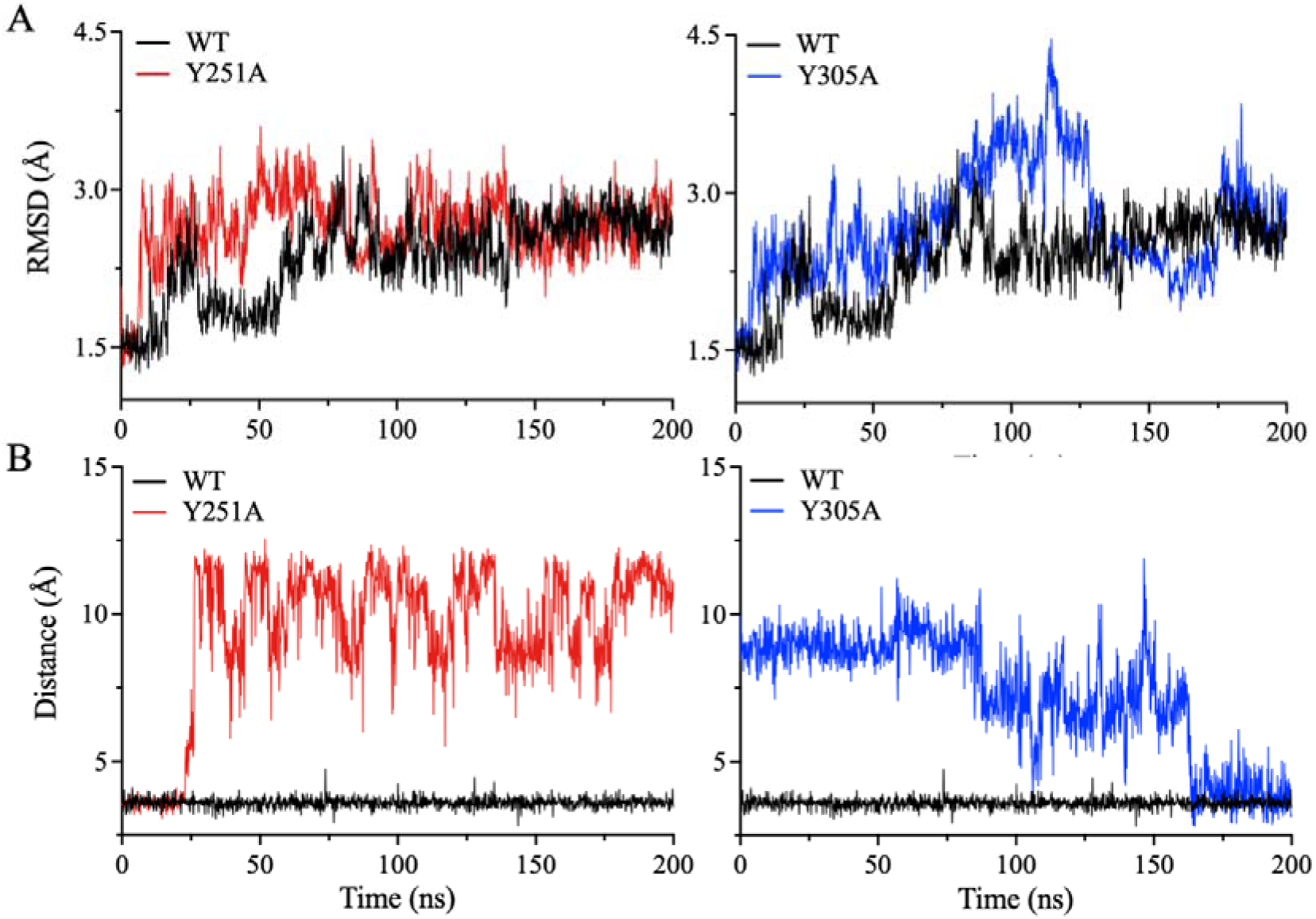
Molecular dynamics simulations of Pocket 3 mutants. (**A**) The root mean square deviation (RMSD) of the protein backbone was determined for WT PLpro and the Pocket 3 mutants Y251A and Y305A. MD simulations of Pocket 1 and 2 mutants did not reveal major structural variations between the mutants and WT PLpro. **(B)** Time-dependence analyses of the distance between the side chains of Y213 of the finger domain and E307 of the palm domain throughout the 200 ns of MD simulations. The distance between the side chains of Y213 and E307 is 2.5 Å in the crystal structure of WT PLpro.

To identify potential factors contributing to the observed instability of Y251A and Y305A in the palm domain during the MD simulations, the distances between Y251A and Y305A and their surrounding residues were comprehensively analyzed. Consistent with the high dynamics observed during the MD simulations, the distances between PLpro domains increased. The distance between the side chain of Y213 of the fingers domain and the side chain of E307 of the palm domain, an interaction close to the positions of Y251 and Y305, was selected to monitor the destabilization of the interaction distances between the fingers and palm domains (**Figure 7B**). In WT PLpro, the distance between Y213 and E307 was maintained at ∼4.3 Å throughout the 200-ns simulation time. In the Y251A mutant, the distance between Y213 and E307 remained at ∼4.3 Å for the first 25 ns of the simulations and then increased to ∼9.60 Å throughout the subsequent duration of the MD run (**Figure 7B**). By contrast, in the simulations of the Y305A mutant, the distance between Y213 and E307 immediately increased, with an average distance of 10 Å over the first 90 ns. The distance then briefly decreased to ∼7.5 Å for the subsequent 70 ns before stabilizing at 4.5 Å for the remaining 40 ns.

## Discussion

The discovery and development of antiviral therapeutics are of paramount importance in the fight against COVID-19 and the spread of SARS-CoV-2. The processing of viral polyproteins by proteases to produce new viral particles is a conserved step in coronavirus maturation ^24–27^. PLpro is not just the main protease of SARS-CoV-2; it has DUB and deISGylating activities that inhibit host innate immunity responses ^46, 47^. Thus, PLpro is an attractive target for the development of antiviral therapeutics against COVID-19 ^10, 11^.

In this study, we present a novel computational and biochemical approach for discovering allosteric sites on the surface of PLpro. Similar analyses of 3CLpro, the other main protease of SARS-CoV-2, identified novel allosteric sites on the surface of 3CLpro with inhibitory potential^48^. Here, three potential surface pockets on the face of PLpro opposite from the active site were computationally identified (**Figure 1**). Residues in these pockets that facilitate side chain–side chain or side chain–backbone interactions were mutated to biochemically assess their importance for proteolytic activity. Identifying key amino acid contacts in PLpro will aid the screening and identification of small-molecule inhibitors of PLpro that hold potential as antiviral drugs against COVID-19 and related coronavirus diseases.

### Pocket 1: the highest druggability score

Pocket 1 is located at the interface between the thumb and Ubl domains. Eighteen key residues with intra- and inter-domain polar interactions were identified and mutated to alanine. Alanine substitutions at five residues, T10, D12, T54, Y72, and Y83, eliminated or drastically reduced proteolytic activity to less than 12% of that of WT PLpro. Another three mutations, Y56A, E67A, and Y71A, partially inactivated PLpro, as proteolytic activity was 18%–48% of that of WT. Alanine mutations of the other ten residues of Pocket 1 did not significantly affect PLpro activity. The effects of these mutations on activity can be attributed to the loss of intra- and inter- domain interactions. For example, T10, D12, and N15 of the Ubl domain participate in intra-domain interactions, but only T10A and D12A inactivated PLpro. Interestingly, D12E or D12N partially or fully recovered activity, respectively. The lower catalytic activity of D12E might be due to the disruption of bonding interactions with T10 and T15 by the longer side chain of glutamate. D12N may provide the correct side chain length and polar interactions, given the similar sizes of asparagine and aspartate. Another intra-domain interaction in Pocket 1 important for PLpro activity is the side chain interaction between T54 and N13 in the Ubl domain. T54A but not N13A completely inactivated PLpro. This may indicate that the side chain interaction of T54 with N13 is not important for PLpro activity. T54 may also hydrogen bond the backbone carbonyl oxygen of F8 in the Ubl domain, and this interaction might be important for PLpro activity. Overall, these results indicate that intra-domain interactions are important for maintaining the structural fold of the Ubl domain and the activity of PLpro.

The other two residues in Pocket 1 that inactivated PLpro when mutated to alanine are Y72 and Y83, both in the thumb domain. Y72 is within a short hydrogen bond distance of 2.6 Å from the backbone carbonyl oxygen of V11, indicating the importance of inter-domain interactions between the Ubl and thumb domains. While Y72T retained only 6% of WT activity, Y72F fully recovered the proteolytic activity of PLpro. These results show that the large aromatic ring of tyrosine is important for maintaining local interactions that support the proper structural fold of PLpro. Y83 forms an intra-domain interaction with N146 of the thumb domain and an inter-domain interaction with Y56 of the Ubl domain. Introducing N146A did not reduce PLpro activity, indicating that the intra-domain interaction of Y83 with N146 is relatively unimportant for activity. By contrast, introducing Y56A reduced PLpro activity to 44% of that of WT, demonstrating that inter-domain interactions between the Ubl and thumb domains are important for PLpro activity. Introducing Y83F recovered 96% of WT activity, whereas Y83T had only 8% of WT activity. Thus, the hydrophobic interactions of Y83 are more influential than its polar interactions for proper folding and catalysis. The Y56T and Y56F mutants both had >75% of WT activity, indicating that both the creation of a local hydrophobic environment and polar interactions are important for activity.

Overall, four tyrosine residues in Pocket 1—one from the Ubl domain (Y56) and three from the thumb domain (Y71, Y72, and Y83)—play critical roles in maintaining active PLpro by providing structural support for the Ubl domain and its contact points with the thumb domain (Figure S8). The tyrosine cluster stabilizes the interactions at the Ubl–thumb interface. By contrast, intra-domain interactions in the thumb domain do not have a major impact on PLpro enzyme.

### Pocket 2: least potential as an allosteric site

Pocket 2 is the only pocket located at the interface of three domains: thumb–fingers–palm. Only four of the 21 residues in Pocket 2 eliminated or reduced PLpro activity to <16% of WT when mutated to alanine: Q237, S239, H275, and S278. Inter-domain interactions dominated the effects of these residues on PLpro activity. Q237 and S239 interact with one another and are at the boundaries of the fingers and palm domains: Q237 is the last residue of the fingers domain, and S239 is the second residue of the palm domain. S239 is located on a loop that connects the last strand of the three-stranded antiparallel β-sheet of the fingers domain with the first strand of the six-stranded antiparallel β-sheet of the palm domain. The tight hydrogen bonding interactions between Q237 and S239 stabilize the loop between the two sheets of the fingers and palm domains in a coiled tight 180° turn. These interactions are important for maintaining the two domains in conformations that stabilize active PLpro.

H275 of the palm domain forms multiple inter-domain interactions with residues in the thumb domain, most importantly Q122, which is a short hydrogen bond distance of 2.8 Å from H275. Introducing Q122A reduced the activity of PLpro. Another inter-domain interaction in the vicinity of H275 is between Q122 of the thumb domain and T277 of the palm domain, and introducing T277A reduced PLpro activity. Furthermore, compared to WT PLpro, Q122A, T277A, and S278A had lower thermal stabilities and *t*_1/2_ values. Collectively, these findings support the importance of this network of inter-domain interactions at the thumb–palm interface in maintaining the thermal stability and catalytic activity of PLpro.

Although multiple intra-domain interactions in the thumb, fingers, and palm domains are observed in Pocket 2, only those of S278A of the palm domain were essential for PLpro activity. Introducing S278A reduced PLpro activity to 16% of that of WT. S278 participates in intra-domain interactions with the backbones of G256 and K279. H255 also participates in this interaction network and hydrogen bonds with the backbone of K279; introducing H255A reduced PLpro activity to 43% of that of WT. In the thumb domain, intra-domain interactions facilitated by Q133 and R140 are important for PLpro activity. Interestingly, most of the mutations in Pocket 2 reduced *K*_m_, indicating a higher affinity for the peptide substrate. However, these increases in *K*_m_ were accompanied by decreases in *k_cat_*, i.e., catalytic efficiency.

### Pocket 3: A high number of amino acids inactivated the PLpro protease

Of the three pockets, Pocket 3, which is located at the finger–palm interface, contains the largest number of residues that inactivated PLpro when mutated to alanine. Thirteen key residues that facilitate intra- and inter-domain polar interactions were identified in Pocket 3. Alanine substitutions at five residues, S212A, Y213A, Y251A, K254A, and Y305A, inactivated PLpro. Another five mutations—E214A, K217A, T259A, K306A, and E307A— partially inactivated PLpro, with relative activities of 21% to 53% compared to WT. The remaining three residues of Pocket 3 do not significantly affect PLpro activity.

Most of the residues in Pocket 3 that are essential for PLpro activity participate in inter-domain interactions between the fingers and palm domains. The inter-domain interaction between the side chains of S212 and Y251 is crucial for PLpro activity, and both S212A and Y251 were classified as inactive. *k*_cat_ was only measurable for S212A and was ∼10-fold lower than that of WT. Interestingly, S212T failed to restore activity, possibly because the side chain of threonine is bulkier than that of serine. In addition, the *t*_1/2_ of S212A was ∼3-fold shorter than that of WT, further highlighting the significance of this residue for the structural and functional stability of PLpro. A second set of inter-domain interactions occurs between Y213 of the fingers domain and Y305 and E307 of the palm domain. Y213A and Y305A were inactive, whereas E307A had a relative activity of 29%. Although Y213T was inactive, introducing Y213F partially recovered proteolytic activity to 28% of that of WT, indicating that the hydrophobic interactions of the side chain of Y213 are important for activity. By contrast, neither Y305F nor Y305T were active, demonstrating that both the large hydrophobic aromatic ring and hydroxyl group of tyrosine are important. The third inter-domain interaction is between the side chain of K306 of the palm domain and the backbone carbonyl oxygen of Q122 of the thumb domain. Alanine mutations at these positions reduced but did not eliminate PLpro activity.

In Pocket 3, intra-domain interactions were observed between E214 and K217 of the fingers domain and between E252, K254, T257, T259, E295, and K306 of the palm domain. Only the alanine mutation at K254 drastically reduced PLpro activity, and the K254R mutant had 67% relative activity. Alanine mutations of the other residues reduced the catalytic activity of PLpro, with relative activity values of 44% to 76%. These results suggest that inter-domain interactions in Pocket 3 contribute more than intra-domain interactions to the activity and stability of PLpro.

### Kinetic and dynamic stabilities are important for an active PLpro

The DSC analysis showed that the inactive mutants had higher melting points and lower enthalpies of unfolding than WT PLpro. These mutations likely altered the structural fold of PLpro, as the magnitude of Δ*H*_cal_ is correlated with the extent of bonding interactions in a protein’s structure. Positive values of Δ*H*_cal_ indicate that hydrophilic interactions are dominant, whereas negative values indicate that hydrophobic interactions dominate the structure ^49^. Thus, the lower Δ*H*_cal_ values of the inactive mutants indicate that they were more hydrophobic than WT PLpro, despite greater thermodynamic stability. Moreover, the greater thermodynamic stability of the inactive mutants indicates that their inactivity was due to structural changes preventing catalysis and not to destabilization of the structural fold.

The MD simulations of the inactive mutants Y251A and Y305A of the palm domain in Pocket 3 demonstrated that inter-domain interactions of these residues are important for maintaining the distance between the palm and fingers domains. This distance was monitored by measuring the distance between the side chains of Y213 and E307 during the MD simulations of Y251A and Y305A mutants and comparing it with the WT enzyme. The Y213–E307 distance increased in Y251A and Y305A compared with WT during the MD simulations, which further highlights the importance of Y251 and Y305 in stabilizing the inter-domain interactions between the fingers and palm domains for active PLpro. The interactions facilitated by Y251 and Y305 are essential for preserving the functional fold and structural stability of PLpro. Hence, the loss of activity in the Y251A and Y305A mutants can be partially attributed to increased distances between the finger and palm domains of PLpro. These findings underscore the complexity and critical role of inter-domain interactions in maintaining the structural and functional integrity of PLpro.

## Conclusion

Each of the computationally identified pockets was assigned a *D*score based on the cavity size, enclosure, and hydrophobicity of each pocket. Pockets 2 and 3 have similar druggability scores, sizes, and enclosures, but Pocket 3 has higher hydrophobicity. Of the 52 residues mutated in this study, 14 reduced relative activity (compared to WT) to less than 15% when mutated to alanine, and another 16 reduced relative activity to less than 60% when mutated to alanine. Although Pocket 1 has the highest *D*score (i.e., greatest potential druggability), Pocket 3 is the most promising allosteric pocket because it contains the largest number of residues that are required for PLpro activity. The biochemical analyses in this study provide important information on the types of interactions that should be disrupted by inhibitors, which is important for guiding the computational screening of small-molecule libraries and the design of antiviral drugs targeting SARS-CoV-2 PLpro.

## Materials and Methods

### Druggability assessment and identification of surface pockets on SARS-CoV-2 PLpro

Twenty-two crystal structures of SARS-CoV-2 PLpro were downloaded from the Protein Data Bank (http://www.rcsb.org). All structures were of proteins without mutations that were solved by X-ray crystallography. MOE (Molecular Operating Environment, 2021) was used to remove solvent atoms from each crystal structure and add any missing atoms, residues, chains, or loops using the protein preparation module. Next, Protonate3D was used to assign each atom a unique protonation state. The prepared crystal structures were then imported into Maestro (Schrödinger, 2021), where the Protein Preparation Wizard (Schrödinger, 2021) module was used to refine the structures and to maintain structural integrity. Hydrogens were added using hydrogen bond optimization, and the structures were subsequently subjected to restrained minimization to achieve a reduced energy state.

Pockets were identified using the SiteMap (Schrödinger, 2021) module with default settings. To prevent any bias from utilizing ligands or peptides to identify pockets, the "Identify top-ranked possible receptor binding sites" option was used. SiteMap computes numerous physiochemical parameters for each pocket, including size, exposure, hydrophobicity, hydrophilicity, and volume^50^. It then assigns a *D*score to each pocket using a weighted equation (Eq. 1):

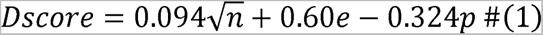

where n is the number of site points, e is the enclosure factor, and p denotes hydrophilicity.

### Enzyme titration and initial velocity studies of wild-type (WT) and mutant PLpro

The gene encoding recombinant WT or mutant PLpro with an N-terminal Hisx6 tag was introduced into the bacterial expression vector pET-28b (+) by GenScript, Inc. (Piscataway, NJ). The resulting plasmid was used to transform *E. coli* BL21-CodonPlus-RIL (Stratagene) for protein expression, which was induced as described previously (Shetler, Ferreira *et al.* 2022). The bacterial cell pellets were homogenized in lysis buffer [25 mM Tris pH 7.5, 150 mM NaCl, 5 mM imidazole, 3 mM β-mercaptoethanol (βME), and 0.1% protein inhibitor cocktail (Sigma-Aldrich: P8849)], and after centrifugation, the supernatant was loaded onto a nickel column pre-equilibrated with binding buffer (25 mM Tris, 150 mM NaCl, 5 mM imidazole, and 3 mM βME). The column was washed with binding buffer, and protein was eluted with binding buffer supplemented with 300 mM imidazole. The fractions from the nickel column were loaded onto a HiLoad Superdex 200pg (16/600) size-exclusion chromatography column (GE Healthcare) on an ÄKTA pure 25 chromatography system (Cytiva, USA). The column was pre-equilibrated with 20 mM HEPES pH 7.5, 150 mM NaCl, and 0.5 mM tris(2-carboxyethyl)phosphine (TCEP). The PLpro-containing fractions were collected and concentrated to ∼150 μM. The protein concentration and purity were determined by the Bradford assay ^51^ and SDS-PAGE, respectively.

PLpro enzyme titration and initial velocity analyses were performed using a Cytation 5 multi-mode microplate reader (BIOTEK Instruments, Winooski, VT) as described previously ^15^. The peptide substrate contained the PLpro cleavage site Leu-Arg-Gly-Gly flanked by a fluorescent AMC (7-amino-4-methyl coumarin) tag and corresponding quencher CBZ (carbobenzoxy) (*i.e.*, CBZ-LRGG-AMC). The proteolytic reaction was initiated by adding PLpro to the peptide substrate in buffer (20 mM HEPES pH 7.5, 150 mM NaCl, 1 mM TCEP, and 1 mM EDTA) containing 2% DMSO (v/v) to enhance the solubility and solubility of the peptide ^15^. The PLpro reaction was monitored in a 96-well microplate at 25 °C for 10 min at excitation and emission wavelengths of 340 nm and 487 nm, respectively. Cleavage of the peptide substrate by PLpro increased the AMC fluorescent signal.

Enzyme titration analyses were performed to assess the effect of a single amino acid substitution on PLpro proteolysis. In brief, the enzymatic activity of WT or mutant PLpro was assayed at different enzyme concentrations ranging from 0.5 to 5.0 μM at a fixed peptide substrate concentration of 200 μM. Next, kinetic parameters, including *K*_m_ and *k_cat_*, were determined by performing initial velocity studies of the WT enzyme and the catalytically active mutants. The concentration of the peptide substrate was varied from 20 to 800 μM at a fixed concentration of PLpro. The cleavage rate data were fit to the Michaelis–Menten equation using the global fitting analysis function in the kinetics module of SigmaPlot (Systat Software, Inc). Error bars were calculated from triplicate measurements of each reaction, and the results are presented as the mean ± SD.

### Differential scanning calorimetry (DSC) and thermal inactivation kinetics of PLpro variants

The thermodynamic stability of WT and mutant PLpro was measured using a Nano DSC instrument (TA Instruments) calibrated using chicken egg white lysozyme, an external standard in the TA Instruments test kit (602198.901). The DSC thermograms were acquired at a PLpro concentration of 0.3 mg/mL in 20 mM HEPES pH 7.0, 150 mM NaCl, and 0.5 mM TCEP. The protein samples were heated from 15 °C to 75 °C at a scan rate of 1 °C/min and 3 atm pressure. Background scans were obtained by loading degassed buffer in both the reference and sample cells and heating at the same rate. The DSC thermograms were corrected by subtracting the corresponding buffer baseline and converted to plots of excess heat capacity (*C*_p_) as a function of temperature. The sample melting point (*T*_m_) was determined at the maximum temperature of the thermal transition, and the calorimetric enthalpy (Δ*H_cal_*) of the transition was estimated from the area under the thermal transition using Nano Analyzer software (*TA* Instruments).

To determine the thermal inactivation kinetics of the PLpro mutants, proteolytic activity was assayed after incubating the enzyme at 37 °C for different time intervals. The enzymatic activity of PLpro was determined as described above, and the decay in enzymatic activity was fit to a one-phase decay equation (Eq. 2) using Prism 9 (GraphPad Software):

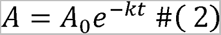

where *A* is the relative activity of the enzyme at incubation time *t*, *A*_0_ is the activity of the enzyme at *t* = 0, and *k* is the rate constant for enzyme inactivation. The half-life (*t*_1/2_) was defined as the time at which enzymatic activity was reduced to half and calculated as *t*_1/2_ = *ln*(2) /*k*.

### Molecular dynamics (MD) simulations of WT and mutant PLpro

The residue scan module in MOE (Chemical Computing Group) was used to introduce single amino acid mutations into the structure of PLpro, including the inactivating mutations Y251A and Y305A. MD simulations of the WT and mutant enzymes were performed using the AMBER18 package ^52^. The ff19SB force field was used to assign partial charges and other parameters to the protein. The protein system was constructed using the xleap module of AmberTools. Subsequently, neutralization was achieved by introducing Na^+^ counter ions, and solvation was performed in a truncated octahedral box of TIP3P water. The energy of the system underwent a two-step minimization process using the PMEMD program in the AMBER18 package ^52^. First, all solute atoms were restrained with a force constant of 500 kcal mol^-1^ Å^-2^. This was followed by a minimization step without any constraints. Under the NVT ensemble, the system was gradually heated from 0 to 300 K. The SHAKE algorithm was employed for all hydrogen atoms that form bonds with a collision frequency of 1.0 ps^-1^. Following this, a 200-ns simulation of the protein was conducted in the NPT ensemble, where the system’s temperature and pressure were maintained at 300 K and 1.01 x 10LJ Pa, respectively. All trajectories were analyzed using DBSCAN ^53^ in the CCPTRAJ module in AmberTools ^54^. Cluster analysis was conducted on every tenth frame (10 ns), excluding ions and solvent molecules from the system. The distance cutoff (ε) for cluster formation was maintained at the default value of 3.0. The CCPTRAJ tool was used to analyze the trajectory files by generating an RMSD plot.

