## Supplementary data for "Identification of Novel Allosteric Sites of SARS-CoV-2 Papain-Like Protease (PLpro) for the Development of COVID-19 Antivirals"

**Table S1.** Mean values of the druggability score (*D*score) and other SiteMap physiochemical parameters obtained from the 22 PLpro structures analyzed in this work.

| **Site** | ***D*score** | **Size** | **Enclosure** | **Hydrophilicity** | |
| --- | --- | --- | --- | --- | --- |
| **Active Site** | 0.88 | 75 | 0.68 | | 0.99 |
| **Pocket 1** | 0.75 | 61 | 0.67 | | 1.20 |
| **Pocket 2** | 0.57 | 29 | 0.58 | | 0.89 |
| **Pocket 3** | 0.57 | 31 | 0.62 | | 0.99 |

**Table S2.** Residues of the computationally predicted druggable pockets of SARS-CoV-2 PLpro. Only amino acids with potentially important side chain interactions were selected for further mutagenesis and biochemical analyses.

| **Site** | **Residues** |
| --- | --- |
| **Pocket 1** | D12, N13, I14, L36, D37, G38, K53, T54, F55, Y56, Y71, Y72, Y83, L87, K91, A131, D134, A135, R138, E143, A144, A145, N146, L150 |
| **Pocket 2** | L120, Q122, I123, E124, L125, K126, F127, Q133, Y136, Y137, R140, N177, L178, S180, C181, E238, S239, P240, F241, G256, F258, T259, T277, S278, K279 |
| **Pocket 3** | S212, S213, E214, K217, Y251, E252, L253, K254, T257, F258, V303, Y305, K306, E307 |

**Table S3.** Kinetic parameters of Pocket 1 mutants of PLpro.

| **Variant** | ***k_cat_* (min^−1^)** | ***K*_M_ (µM)** | ***k_cat_*/*K*_M_ (min^−1^ • µM^−1^)** |
| --- | --- | --- | --- |
|  | Fold Change | Fold Change | Fold Change |
| WT | 4.75 ± 0.08 | 231 ± 6 | 0.0206 ± 0.0003 |
| D12A | 0.44 ± 0.01 | 273 ± 2 | 0.0016 ± 0.0001 |
|  | **- 10.8** | **+ 1.2** | **- 12.8** |
| D12E | 0.81 ± 0.03 | 400 ± 10 | 0.0020 ± 0.0001 |
|  | **-5.9** | **+ 1.7** | **- 10.2** |
| D12N | 2.80 ± 0.07 | 420 ± 40 | 0.0067 ± 0.0003 |
|  | **-1.7** | **+ 1.8** | **- 3.1** |
| H17A | 2.80 ± 0.10 | 200 ± 10 | 0.0140 ± 0.0002 |
|  | **-1.7** | **- 1.2** | **- 1.5** |
| Y56A | 1.70 ± 0.20 | 300 ± 20 | 0.0058 ± 0.0003 |
|  | **-2.8** | **+ 1.3** | **- 3.6** |
| Y56F | 2.77 ± 0.07 | 400 ± 10 | 0.0070 ± 0.0001 |
|  | **- 1.7** | **+ 1.7** | **-2.9** |
| Y56T | 2.10 ± 0.20 | 470 ± 40 | 0.0045 ± 0.0003 |
|  | **- 2.3** | **+ 2.0** | **- 4.6** |
| E67A | 0.70 ± 0.06 | 290 ± 20 | 0.0024 ± 0.0001 |
|  | **- 6.8** | **+ 1.3** | **- 8.6** |
| Y71A | 2.29 ± 0.05 | 137 ± 3 | 0.0167 ± 0.0001 |
|  | **- 2.1** | **- 1.7** | **- 1.2** |
| Y71F | 2.90 ± 0.70 | 700 ± 80 | 0.0042 ± 0.0007 |
|  | **- 1.6** | **+ 3.0** | **- 4.9** |
| Y71T | 1.90 ± 0.10 | 460 ± 30 | 0.0042 ± 0.0002 |
|  | **- 2.5** | **+ 2.0** | **- 4.9** |
| Y72F | 2.30 ± 0.07 | 420 ± 60 | 0.0056 ± 0.0006 |
|  | **- 2.1** | **+ 1.8** | **- 3.7** |
| Y72T | 0.22 ± 0.01 | 420 ± 30 | 0.0005 ± 0.0001 |
|  | **- 22.0** | **+ 1.8** | **- 40.2** |
| Y83A | 0.55 ± 0.03 | 206 ± 5 | 0.0027 ± 0.0001 |
|  | **- 8.6** | **- 1.1** | **- 7.7** |
| Y83F | 2.20 ± 0.10 | 460 ± 40 | 0.0047 ± 0.0004 |
|  | **- 2.2** | **+ 2.0** | **- 4.3** |
| Y83T | 0.25 ± 0.04 | 440 ± 50 | 0.0006 ± 0.0001 |
|  | **-19** | **+1.9** | **-37.3** |
| K91A | 3.70 ± 0.30 | 210 ± 20 | 0.0175 ± 0.0006 |
|  | **- 1.3** | **- 1.1** | **- 1.2** |
| D134A | 4.83 ± 0.05 | 267 ± 7 | 0.0181 ± 0.0003 |
|  | **+ 1.0** | **+ 1.2** | **-1.1** |
| R138A | 5.20 ± 0.60 | 230 ± 30 | 0.0221 ± 0.0008 |
|  | **+ 1.1** | **+ 1.0** | **+ 1.1** |
| E143A | 5.20 ± 0.08 | 210 ± 8 | 0.0248 ± 0.0006 |
|  | **+ 1.1** | **- 1.1** | **+ 1.2** |
| N146A | 6.40 ± 0.20 | 240 ± 10 | 0.0268 ± 0.0005 |
|  | **+ 1.3** | **+ 1.0** | **+ 1.3** |

**Table S4.** Kinetic parameters of Pocket 2 mutants of PLpro.

| **Variant** | ***k_cat_* (min^−1^)** | ***K*_M_ (µM)** | ***k_cat_*/*K*_M_ (min^−1^ • µM^−1^)** |
| --- | --- | --- | --- |
|  | Fold Change | Fold Change | Fold Change |
| WT | 4.75 ± 0.08 | 231 ± 6 | 0.0206 ± 0.0003 |
| T119A | 6.6 ± 0.4 | 350 ± 50 | 0.0198 ± 0.0004 |
|  | **+ 1.4** | **+ 1.5** | **- 1.0** |
| Q122A | 1.70 ± 0.20 | 190 ± 30 | 0.0088 ± 0.0002 |
|  | **- 2.9** | **- 1.2** | **- 2.4** |
| Q122E | 1.21 ± 0.08 | 470 ± 10 | 0.0026 ± 0.0002 |
|  | **- 3.9** | **+ 2.1** | **- 8.0** |
| Q133A | 3.20 ± 0.10 | 177 ± 8 | 0.0179 ± 0.0009 |
|  | **- 1.5** | **- 1.3** | **- 1.2** |
| Q133N | 1.50 ± 0.10 | 470 ± 60 | 0.0031 ± 0.0001 |
|  | **- 3.3** | **+ 2.1** | **- 6.6** |
| R140A | 2.52 ± 0.05 | 220 ± 4 | 0.0115 ± 0.0001 |
|  | **- 1.9** | **- 1.1** | **- 1.8** |
| N177A | 3.23 ± 0.02 | 270 ± 10 | 0.0118 ± 0.0004 |
|  | **- 1.5** | **+ 1.2** | **- 1.8** |
| D179A | 4.90 ± 0.50 | 470 ± 60 | 0.0103 ± 0.0003 |
|  | **+ 1.0** | **+ 2.1** | **- 2.0** |
| Q236A | 1.04 ± 0.01 | 330 ± 3 | 0.0032 ± 0.0001 |
|  | **- 4.6** | **+ 1.4** | **- 6.5** |
| E238A | 2.90 ± 0.10 | 370 ± 40 | 0.0081 ± 0.0005 |
|  | **- 1.6** | **+ 1.6** | **- 2.5** |
| E238D | 2.00 ± 0.20 | 410 ± 20 | 0.0049 ± 0.0003 |
|  | **- 2.4** | **+ 1.8** | **- 4.2** |
| E238Q | 2.20 ± 0.10 | 420 ± 40 | 0.0053 ± 0.0005 |
|  | **- 2.2** | **+ 1.8** | **- 3.9** |
| S239A | 0.15 ± 0.01 | 330 ± 30 | 0.0004 ± 0.0001 |
|  | **- 32.6** | **+ 1.4** | **- 46.9** |
| H255A | 1.12 ± 0.02 | 301 ± 1 | 0.0037 ± 0.0001 |
|  | **- 4.3** | **+ 1.3** | **- 5.5** |
| T277A | 1.53 ± 0.06 | 150 ± 10 | 0.0106 ± 0.0004 |
|  | **- 3.1** | **- 1.6** | **- 1.9** |
| S278A | 0.89 ± 0.05 | 157 ± 6 | 0.0057 ± 0.0002 |
|  | **- 5.3** | **- 1.5** | **- 3.6** |
| S278T | 0.17 ± 0.01 | 500 ± 30 | 0.0003 ± 0.00002 |
|  | **- 28.7** | **+ 2.2** | **- 62.7** |
| K279A | 4.00 ± 0.20 | 240 ± 10 | 0.0171 ± 0.0003 |
|  | **- 1.2** | **+ 1.0** | **- 1.2** |

**Table S5.** Kinetic parameters of Pocket 3 mutants of PLpro.

| **Variant** | ***k_cat_* (min^−1^)** | ***K*_M_ (µM)** | ***k_cat_*/*K*_M_ (min^−1^ • µM^−1^)** |
| --- | --- | --- | --- |
|  | Fold Change | Fold Change | Fold Change |
| WT | 4.75 ± 0.08 | 231 ± 6 | 0.0206 ± 0.0003 |
| S212A | 0.50 ± 0.03 | 310 ± 30 | 0.0016 ± 0.0001 |
|  | **- 9.6** | **+ 1.3** | **- 12.7** |
| S212T | 0.04 ± 0.01 | 600 ± 100 | 0.00006 ± 0.00001 |
|  | **- 126.3** | **+ 2.8** | **- 343.5** |
| Y213F | 0.90 ± 0.10 | 430 ± 80 | 0.0020 ± 0.0001 |
|  | **- 5.5** | **+ 1.9** | **- 10.2** |
| E214A | 2.80 ± 0.20 | 430 ± 30 | 0.0064 ± 0.0001 |
|  | **- 1.7** | **+ 1.9** | **- 3.2** |
| K217A | 0.13 ± 0.01 | 440 ± 20 | 0.000295 ± 0.0001 |
|  | **- 36.5** | **+ 1.9** | **- 69.7** |
| Y251A | 0.15 ± 0.01 | 310 ± 30 | 0.0005 ± 0.0001 |
|  | **- 31.9** | **+ 1.3** | **- 42.4** |
| Y251F | 2.80 ± 0.50 | 380 ± 70 | 0.0074 ± 0.0007 |
|  | **- 1.7** | **+ 1.6** | **- 2.8** |
| Y251T | 2.37 ± 0.01 | 330 ± 20 | 0.0073 ± 0.0004 |
|  | **- 2.0** | **+ 1.4** | **- 2.8** |
| E252A | 3.66 ± 0.09 | 273 ± 2 | 0.0134 ± 0.0002 |
|  | **- 1.3** | **+ 1.2** | **- 1.5** |
| K254A | 0.44 ± 0.01 | 224 ± 9 | 0.0020 ± 0.0001 |
|  | **- 10.8** | **- 1.0** | **- 10.5** |
| T259A | 1.17 ± 0.01 | 289 ± 3 | 0.0040 ± 0.0001 |
|  | **- 4.0** | **+ 1.3** | **- 5.0** |
| Y305F | 0.24 ± 0.03 | 800 ± 100 | 0.0003 ± 0.00002 |
|  | **- 19.6** | **+ 3.3** | **- 64.3** |
| Y305T | 0.016 ± 0.002 | 600 ± 100 | 0.00003 ± 0.000003 |
|  | **- 297.1** | **+ 2.6** | **- 755.6** |
| K306A | 2.60 ± 0.20 | 210 ± 20 | 0.0121 ± 0.0001 |
|  | **- 1.9** | **- 1.1** | **- 1.7** |
| K306R | 1.45 ± 0.04 | 453 ± 8 | 0.00312 ± 0.0001 |
|  | **- 3.3** | **+ 2.0** | **- 6.4** |
| E307A | 1.17 ± 0.02 | 289 ± 3 | 0.0041 ± 0.0001 |
|  | **- 4.1** | **+ 1.3** | **- 5.1** |

**Table S6.** Thermodynamic parameters of PLpro variants determined by DSC.

| **Pocket** | **Variant** | ***T*_m1_ (°C)** | ***T*_m2_ (°C)** | ***∆H*_cal_ (kJ/mol)** |
| --- | --- | --- | --- | --- |
|  |  | **Fold Change** | **Fold Change** | **Fold Change** |
|  | WT | 44.4 ± 0.4 | 51.9 ± 0.1 | 1094 ± 73 |
| Pocket 1 | T10A | 49.1 ± 0.5 | 55.5 ± 0.2 | 684 ± 131 |
|  |  | **+1.1** | **+1.07** | **-1.60** |
|  | D12A | 53.0 ± 0.3 | 57.2 ± 0.1 | 348 ± 18 |
|  |  | **+ 1.2** | **+ 1.10** | **- 3.15** |
|  | T54A | 47.4 ± 1.5 | 53.0 ± 1.0 | 604 ± 47 |
|  |  | **+1.1** | **+1.02** | **-1.81** |
|  | Y72A | 47.3 ± 0.4 | 55.9 ± 0.1 | 460 ± 35 |
|  |  | **+1.1** | **+1.1** | **-2.4** |
|  | Y83A | 50.7 ± 0.2 | 56.6 ± 0.1 | 758 ± 35 |
|  |  | **+ 1.1** | **+ 1.1** | **- 1.4** |
| Pocket 2 | Q122A | 45.6 ± 0.9 | 55.3 ± 0.5 | 723 ± 25 |
|  |  | **+ 1.0** | **+ 1.1** | **+ 1.1** |
|  | Q237A | 50.9 ± 0.5 | 57.7 ± 0.4 | 694 ± 52 |
|  |  | **+1.2** | **+1.1** | **-1.6** |
|  | E238A | 46.1 ± 0.4 | 51.7 ± 0.2 | 867 ±34 |
|  |  | **+ 1.0** | **+ 1.0** | **- 1.3** |
|  | S239A | 50.7 ± 0.2 | 55.8 ± 0.3 | 1195 ± 103 |
|  |  | **+ 1.1** | **+ 1.1** | **+ 1.1** |
|  | H275A | 48.3 ± 1.6 | 56.2 ± 0.8 | 717 ± 36 |
|  |  | **+1.1** | **+1.1** | **-1.5** |
|  | T277A | 46.1 ± 0.3 | 51.9 ± 0.1 | 2749 ± 146 |
|  |  | **+ 1.0** | **+ 1.0** | **+ 2.5** |
|  | S278A | 43.7 ± 0.2 | 53.2 ± 0.1 | 618 ± 49 |
|  |  | **- 1.0** | **+ 1.0** | **- 1.8** |
| Pocket 3 | S212A | 48.5 ± 0.5 | 56.2 ± 0.8 | 879 ± 83 |
|  |  | **+ 1.1** | **+ 1.1** | **- 1.2** |
|  | Y213A | 45.2 ± 0.2 | 56.5 ± 0.2 | 1854 ± 30 |
|  |  | **+ 1.0** | **+ 1.1** | **+ 1.7** |
|  | E214A | 49.6 ± 0.5 | 54.2 ± 0.3 | 907 ± 25 |
|  |  | **+1.1** | **+1.0** | **-1.2** |
|  | K217A | 47.0 ± 0.2 | 53.7 ± 0.2 | 588 ± 59 |
|  |  | **+1.1** | **+1.0** | **-1.9** |
|  | Y251A | 49.9 ± 0.8 | 57.0 ± 0.6 | 823 ± 74 |
|  |  | **+1.1** | **+1.1** | **-1.3** |
|  | K254A | 46.1 ± 0.7 | 50.5 ± 0.7 | 597 ± 51 |
|  |  | **+ 1.0** | **- 1.0** | **- 1.8** |
|  | T259A | 50.2 ± 0.4 | 56.7 ± 0.5 | 921 ± 59 |
|  |  | **+ 1.1** | **+ 1.1** | **- 1.2** |
|  | Y305A | 50.5 ± 0.3 | 56.6 ± 0.3 | 778 ± 54 |
|  |  | **+ 1.1** | **+ 1.1** | **- 1.4** |

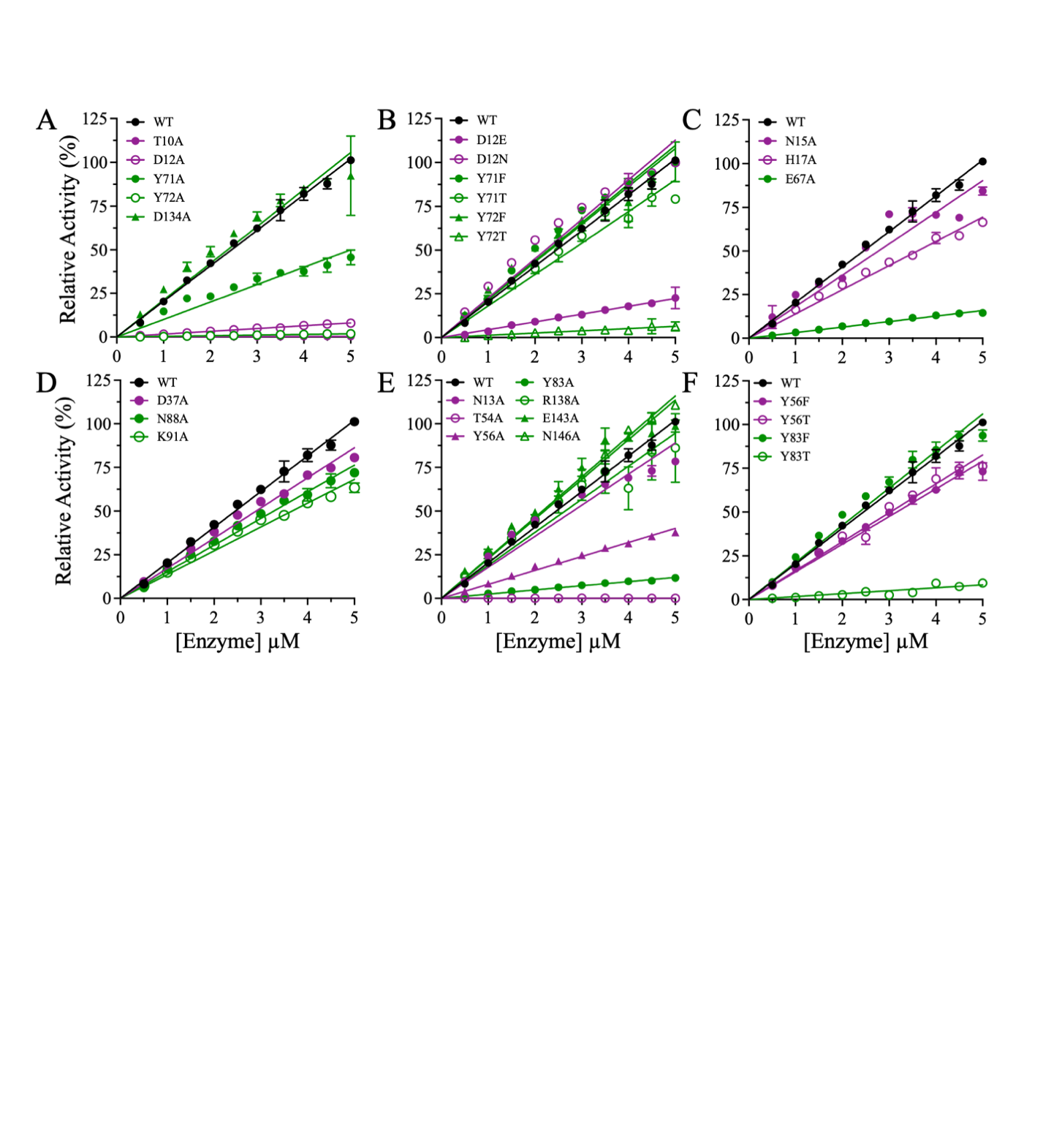

**Figure S1. Relative activities of Pocket 1 mutants of PLpro.** Activity was measured at increasing enzyme concentrations from 0.5 µM to 5.0 μM and a fixed peptide substrate concentration of 200 μM at 25 °C in buffer containing 20 mM HEPES pH 7.5, 150 mM NaCl, 1 mM EDTA, 1 mM TCEP, and 2% DMSO (v/v). The relative activities of the alanine mutants were obtained by normalizing the slopes of the lines to the slope of the line for WT PLpro. WT activity is shown as black-filled circles. The colors of the alanine mutants correspond to their domains: green = thumb and purple = Ubl. The data are presented as the mean ± SD, n = 3.

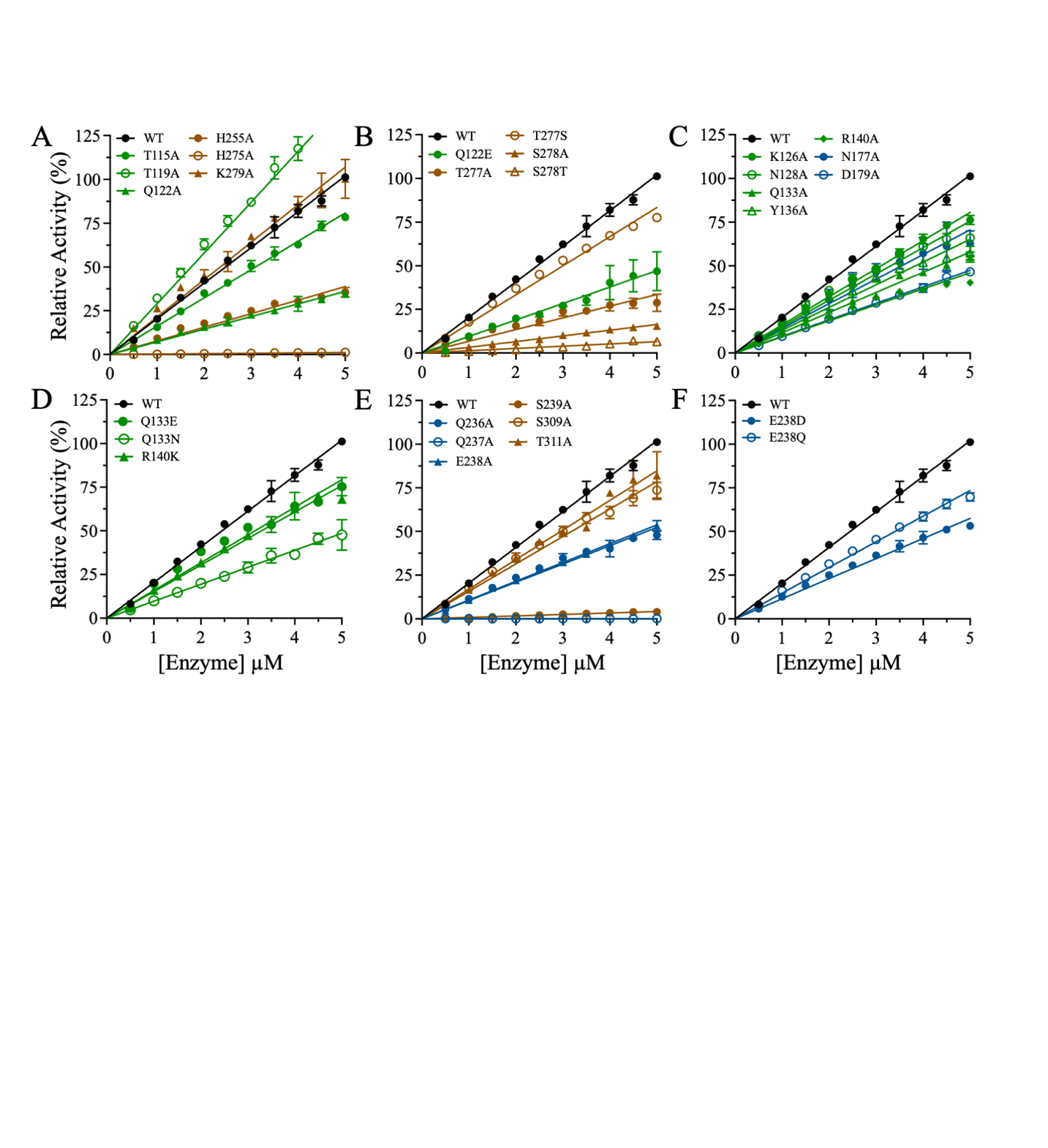

**Figure S2: Relative activities of Pocket 2 mutants of PLpro.** Relative activity was measured as described in Figure S1. WT activity is shown by black-filled circles. The colors of the alanine mutants correspond to their domains: green = thumb, blue = fingers, and brown = palm. The data are presented as the mean ± SD, n = 3.

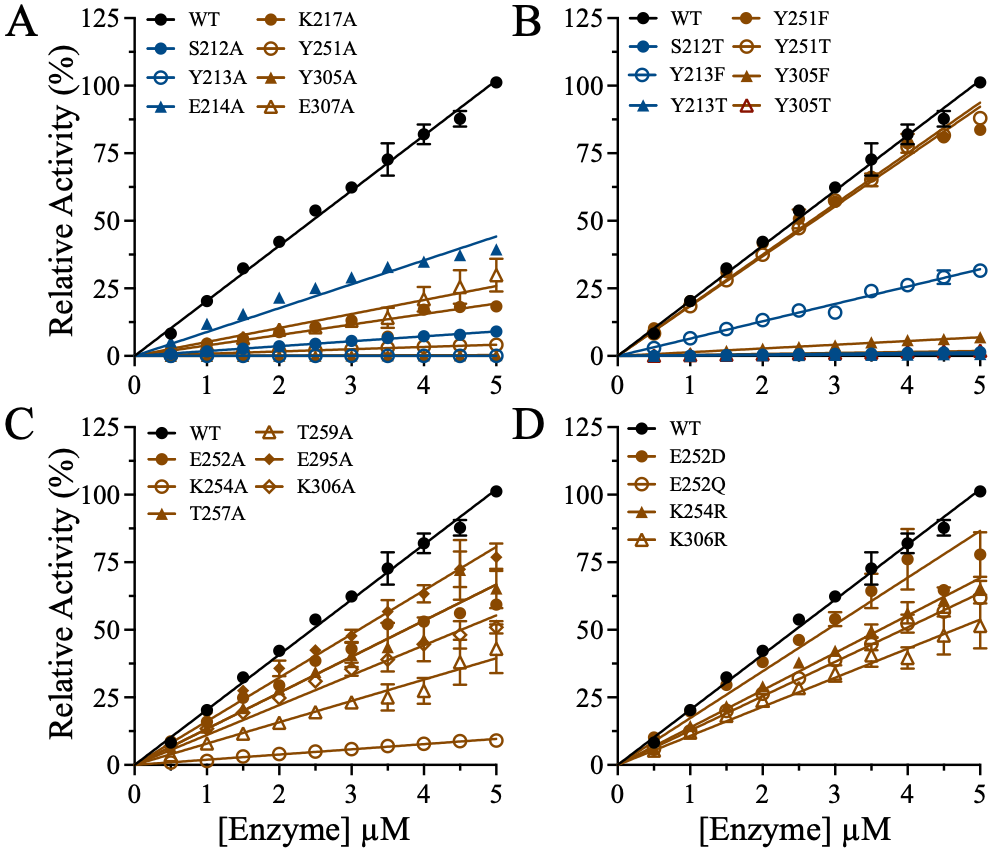

**Figure S3: The relative activities of Pocket 3 mutants of PLpro.** Relative enzymatic activity was measured as described in Figure S1. WT activity is shown by black-filled circles. The colors of the alanine mutants correspond to their domains: blue = fingers and brown = palm. The data are presented as the mean ± SD, n = 3.

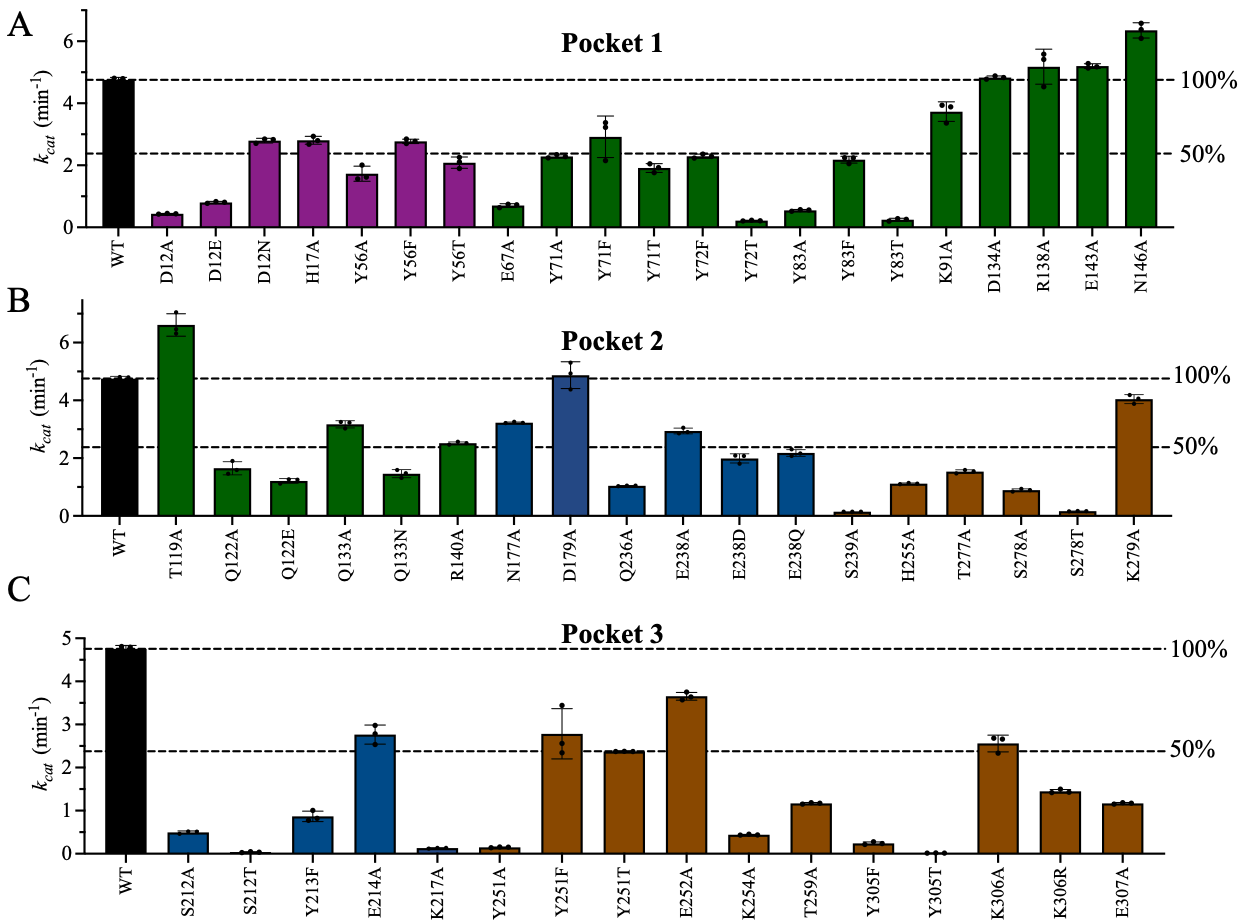

**Figure S4: *k_cat_* values of the enzymatically active PLpro mutants.** Bar plots of the *k_cat_* values of WT PLpro (black bars) and **(A)** Pocket 1 mutants, **(B)** Pocket 2 mutants, and **(C)** Pocket 3 mutants. The initial velocities were measured at 25 °C in buffer containing 20 mM HEPES pH 7.5, 150 mM NaCl, 1 mM EDTA, 1 mM TCEP, and 2% DMSO (v/v). The colors of the alanine mutants correspond to their domains: purple = Ubl, green = thumb, blue = fingers, and brown = palm. The bars represent the mean ± SD, n=3.

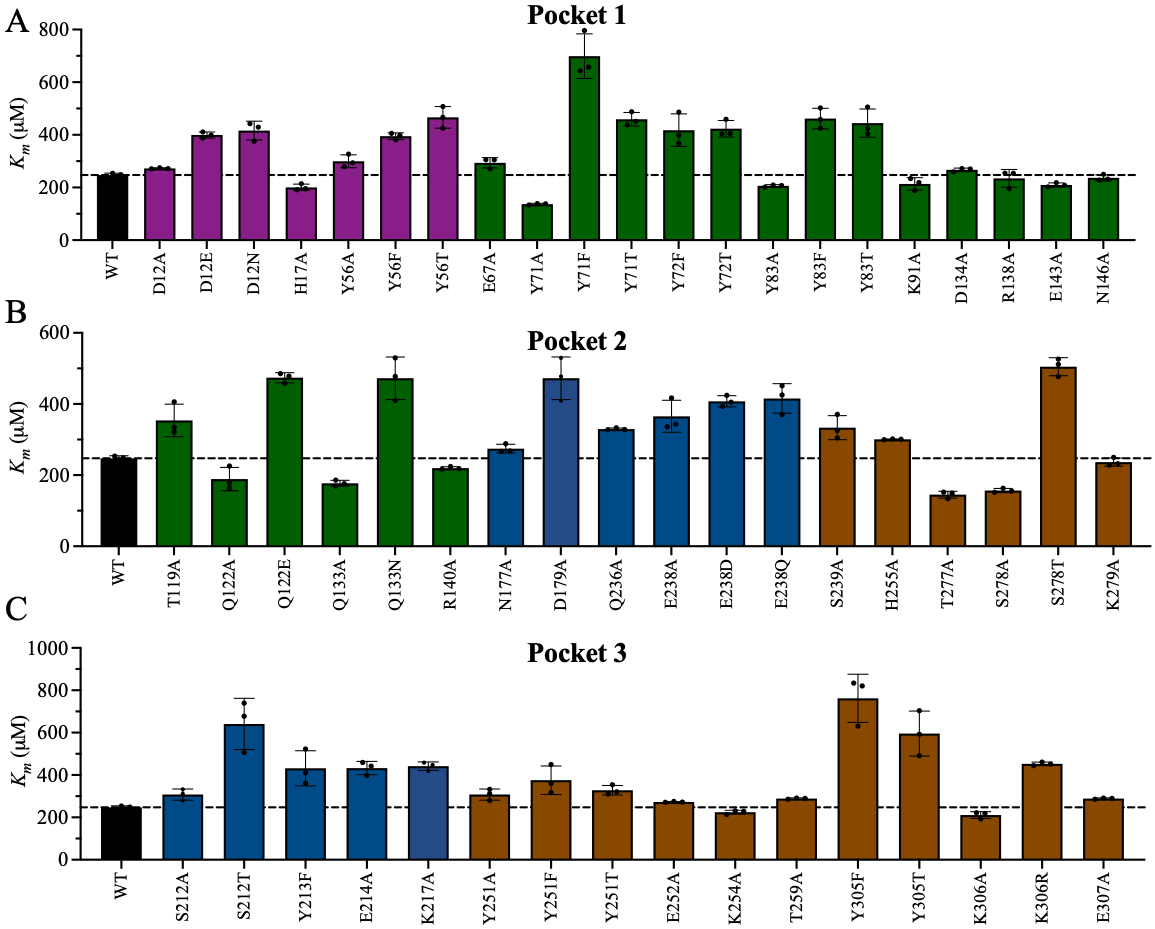

**Figure S5: *K_M_* values of the enzymatically active PLpro mutants.** Bar plots of the *K_m_* values of WT PLpro (black bars) and **(A)** Pocket 1 mutants, **(B)** Pocket 2 mutants, and **(C)** Pocket 3 mutants. The initial velocities were measured as described in Figure S4. The colors of the alanine mutants correspond to their domains: purple = Ubl, green = thumb, blue = fingers, and brown = palm. The bars represent the mean ± SD, n=3.

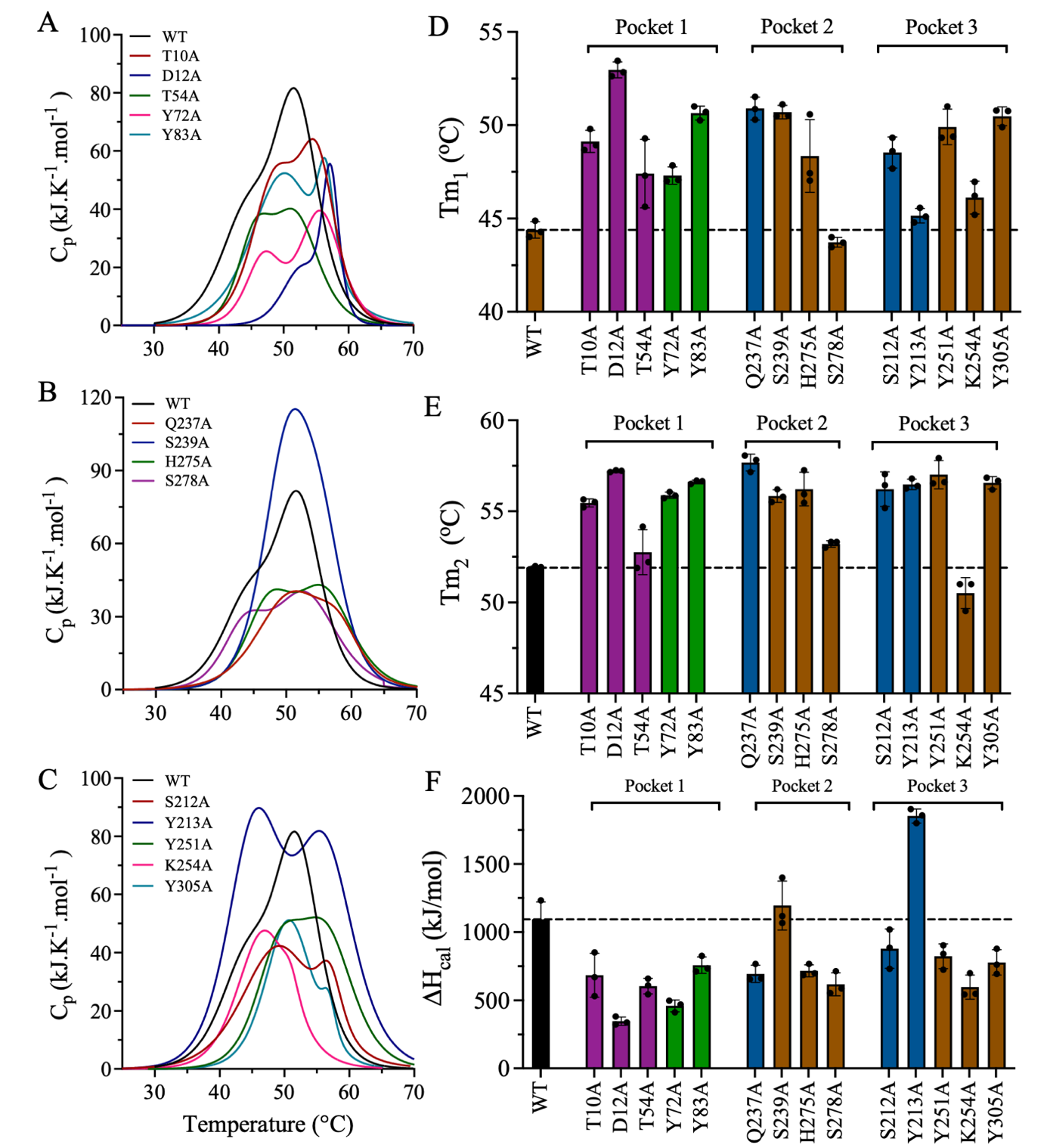

**Figure S6. Thermodynamic parameters of WT PLpro and inactive mutants of Pockets 1–3. (A-C)** DSC thermograms of the inactive mutants of **(A)** Pocket 1, **(B)** Pocket 2, and **(C)** Pocket 3. The thermograms were deconvoluted to a two-state transition with two *T*_m_ values calculated at the two apexes of the thermographic peaks. **(F)** Bar plots of T_m1_ and **(G)** *T*_m2_ of WT PLpro and the inactive mutants of Pockets 1–3. **(H)** Bar plots of the *∆H*_cal_ values of WT PLpro and the inactive mutants of Pockets 1–3 calculated from the area under the thermographic peak. Data are the mean ± SD, n=3.

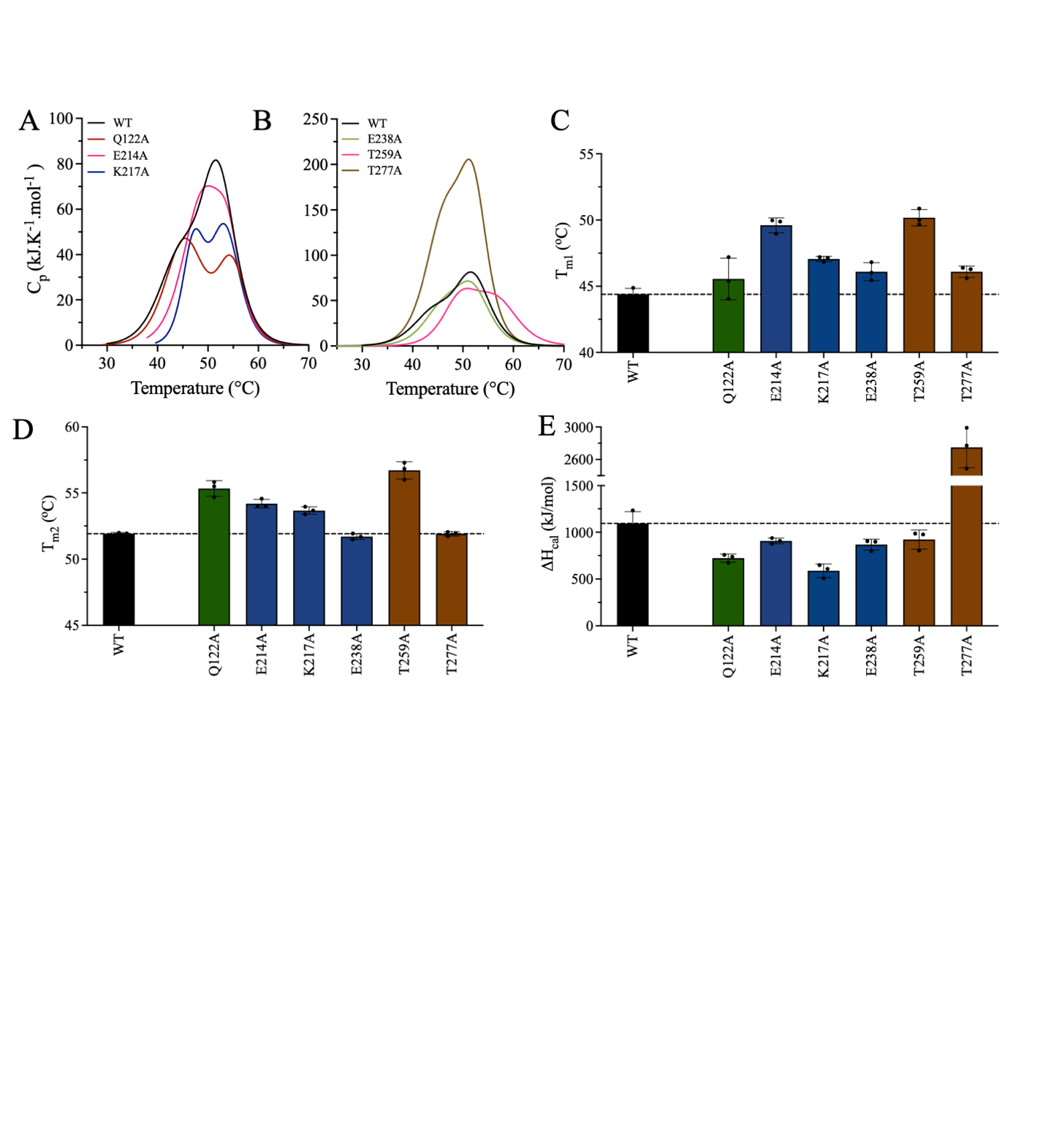

**Figure S7. Thermodynamic parameters of WT PLpro and partially active mutants of Pockets 1–3. (A**–**B)** DSC thermograms of the partially active mutants of Pockets 1–3. **(C)** Bar plots of T_m1_, **(D)** T_m2_, and **(E)** the ∆H_cal_ values of WT PLpro and the partially active mutants of Pockets 1–3. Data are the mean ± SD, n=3.

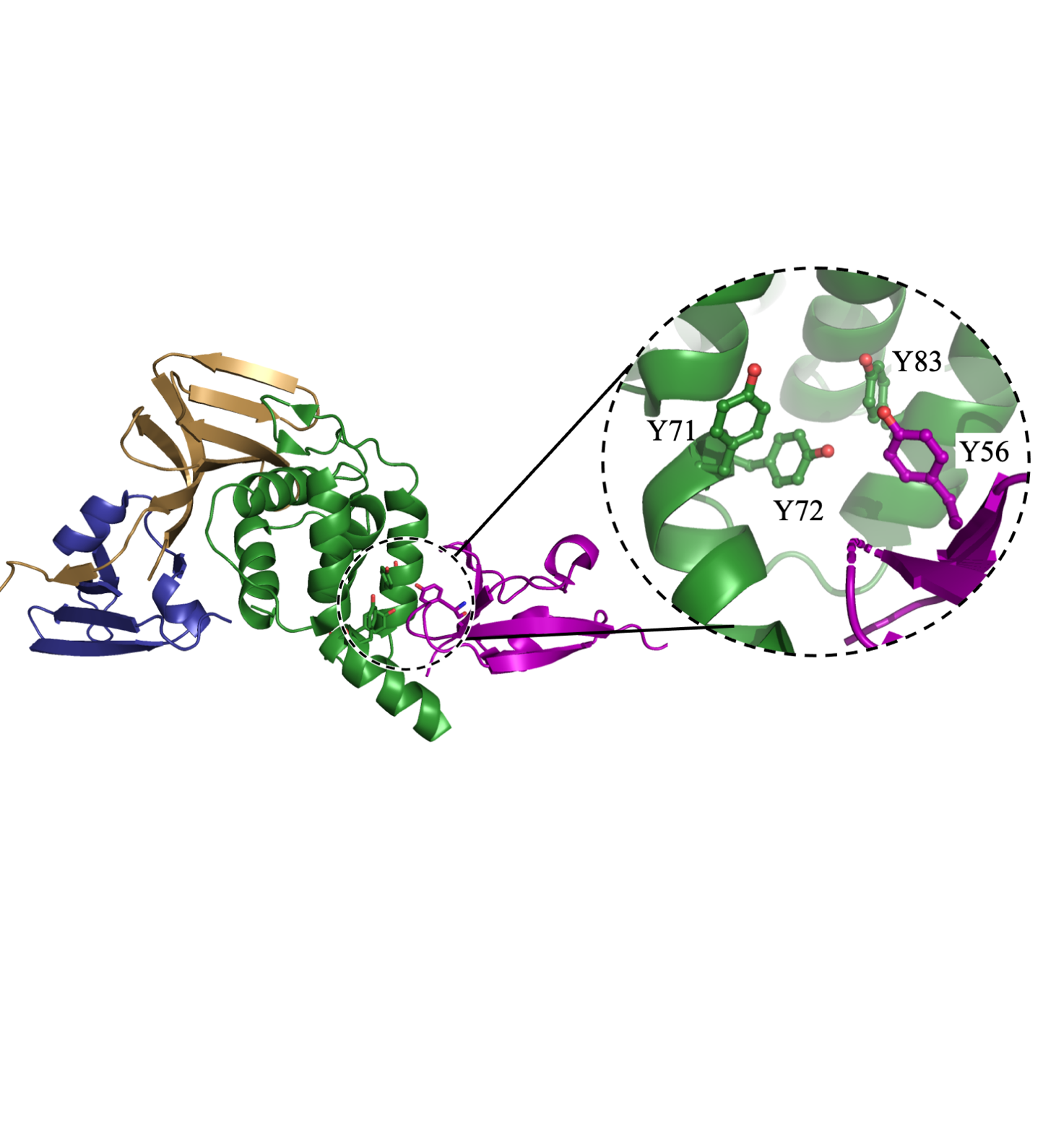

**Figure S8: Cartoon representation of the interface interactions between the Ubl and thumb domains.** Y56 (purple) of the Ubl domain forms a cluster with Y71, Y72, and Y83 (green) of the thumb domain. The tyrosine cluster is thought to stabilize the Ubl–thumb domain interface, which provides structural support for active PLpro. The figure was produced using PyMOL Molecular Graphics System version 2.5.5 (Schrödinger LLC).
